# Muscle power output reflects elevated viscosity in the propulsion system of flying miniature wasps

**DOI:** 10.1101/2025.08.07.668836

**Authors:** Amir Sarig, Thomas Engels, Fritz-Olaf Lehmann, Gal Ribak

**Affiliations:** School of Zoology, Faculty of Life Sciences, Tel Aviv University, Tel Aviv 6997801, Israel; CNRS and Aix-Marseille Université, UMR 7287, Institut des Sciences du Mouvement Etienne-Jules Marey, Marseille, France; Department of Animal Physiology, Institute of Biosciences, University of Rostock, Albert-Einstein-Str. 3, 18059 Rostock, Germany; The Steinhardt Museum of Natural History, Tel Aviv 6997801, Israel

**Keywords:** Insect flight, unsteady aerodynamics, viscous flow, Reynolds number, Computational fluid dynamics, advance ratio, pitch stability, muscle mechanical power, *Eretmocerus mundus*

## Abstract

Air viscosity compromises aerodynamic lift production in the smallest flying insects, leading to increased flight costs. Miniature insects thus utilize both lift and drag for weight support but the exact energetic costs of wing flapping at low Reynolds number are widely unexplored. We estimated flight power in the miniature wasp *Eretmocerus mundus*. Wing kinematics was three-dimensionally reconstructed using high-speed video and computational fluid dynamics simulated air flows, aerodynamic forces and moments. We found an asymmetrical crescent-shaped wingtip trajectory with the upstroke posterior to the downstroke path. This fore-aft distance increases with increasing horizontal flight velocity, maintaining the wing’s backwards rowing motion needed for drag-based propulsion. Although the wing’s lift-to-drag ratio is below unity, lift is the predominant force responsible for weight support and forward thrust. Elevated drag leads to mass-specific mechanical power output of 118±9.0 Wkg^-1^ flight muscle, which exceeds most power estimates reported for other insects, birds and bats. The elevated energetic costs for flight may have fostered the development of bristled wings in miniature insects. Altogether, our study of wingbeat control and flight costs in a miniature insect extends the scope of flight mechanisms to the smallest flying animals revealing limits of miniaturization during the evolution of flight.

## 1. Introduction

Miniature insects with a body length of less than a millimetre must cope with the physical constraints on flight force production at low Reynolds number (Re) below approximately 20 [1–3]. Low Reynolds numbers favour viscous laminar fluid flows in translating and root-flapping wings, attenuating lift production while increasing total drag on the body and its appendages. Thus, viscous drag makes flight of miniature insects energetically more costly compared to flight of their larger relatives [4–6]. Intermediate-sized insects such as fruit flies (Re∼140) may augment lift production using unsteady aerodynamic mechanisms, such as leading-edge vortices, rotational circulation, and wake capture [7–9]. In the viscosity-dominated flight regime of miniature insects, by contrast, the contribution of these mechanisms for lift enhancement is less clear [3, 10, 11]. Recent data, however, show that model wings that flap at Re<5 still produce pronounced lift-enhancing leading-edge vortices [12]. Moreover, many small insect species enhance lift production by clap-and-fling wing kinematics that reinforce the development of aerodynamic circulation at the beginning of the downstroke [13–20]. This augmented lift comes at the cost of elevated drag required to fling the wings apart, which has questioned the energetic advantage of this mechanism for flight of insects [18, 21].

Recently published studies revealed pronounced differences in wing-flapping kinematics of several miniature insect species compared to larger species. During hovering flight, miniature insects typically combine lift- and drag-based propulsion within a complete stroke cycle [22, 23]. This holds for miniature wasps, flies and thrips flapping their wings in a U-shaped, and miniature beetles in a figure-of-eight-shaped, trajectory [23]. In all cases, the wing tip trajectory includes one or two translational rowing phases in which the wings move at elevated angle-of-attack. The first rowing phase occurs during or shortly after the dorsal fling manoeuvre, turning disadvantageous drag into beneficial vertical lift and forward thrust [18]. The second rowing phase may occur at the beginning of the upstroke shortly after the ventral stroke reversal [10]. In addition, an insect must alter flapping kinematics to control manoeuvres, flight velocities and body posture. Larger insect species typically control thrust by changes in body pitch angle, which alters the inclination of the stroke plane and redirects the lift-thrust vector [24–27]. Alternatively, an insect may change the wing’s angle-of-attack during the upstroke that generates thrust owing to a backward rowing movement [28]. However, it is widely unresolved which strategy applies to miniature insects because of their complex mechanisms for the production of aerodynamic forces and moments.

To determine kinematics and energetic costs of wing flapping in miniature insects, this study investigates flight of the tiny parasitic wasp *Eretmocerus mundus* (Hymenoptera: Aphelinidae; [29]). We reconstruct wing and body motion of wasps flying freely at various forward velocities using high-speed videography and derive the appropriate Euler angles. From these data we compute air flows, aerodynamic forces, moments, and power requirements for flight by computational fluid dynamics (CFD). We will show that the low Reynolds number provokes elevated muscle mechanical power output for wing flapping and impose unusual kinematic strategies for body posture and thrust control. In sum, our results eventually reflect the constraints set by drag-based aerodynamic propulsion mechanisms in flying miniature insects.

## 2. Materials and methods

### 2.1. Animals and models

Adults of the parasitic wasp *Eretmocerus mundus* were collected using an aspirator from a population kept in a greenhouse at Tel Aviv University (Fig. 1). The population was maintained in fine mesh cages containing plants infected by *Bemisia tabaci* eggs and larvae as hosts for development. Details on rearing conditions of the host population can be found elsewhere [30]. The collected wasps were transferred to the laboratory in glass vials and their flight behaviour recorded on the day of collection.

**Figure 1.**
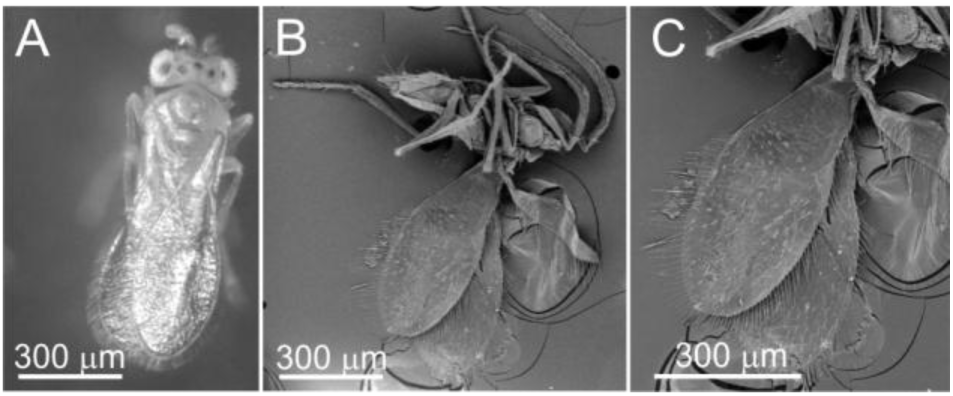
The miniature wasp *Eretmocerus mundus*. (*A*) Top view on an intact animal. (*B*) Scanning electron microscope image showing body, fore- and hindwings with wing bristles. (*C*) Close-up of the forewings.

For CFD analyses, we generated numerical body and wing models based on the morphology of *E. mundus* as described in the Results section. The shape of the forewing and hindwing were derived by outlining their planform contours and then adding the bristles that surround the wing’s edges (electronic supplementary material S1). For simplicity, we used a generic insect body geometry for the simulations. This model was previously constructed [31] but scaled according to *E. mundus* body length and thorax width for the current study. Body mass of the tested animals was estimated from their volume adapting a method previously published for the similarly-sized species *Bemisia tabaci* [30]: from images taken under a dissection microscope we measured length, maximum width and height of the animals and subsequently calculated their volume as an ellipsoid. The resulting volume was multiplied by body density of 700 kgm^-3^ as suggested for the miniature wasp *Anagyrus pseudococci* [32]. This latter value is intermediate relative to values ranging from 400 to 1050 kgm^-3^ reported for other insect species [33].

### 2.2. High-speed video recording and image digitizing

We used three high-speed cameras (Fastcam SA3, Photron Inc.) equipped with 85 mm lenses (Nikkor, Nikon) for recording wing kinematics at 6,000 frames per second. Video images were spatially calibrated with MATLAB script ’DLTcal5’ and by moving an insect mounting pin to known positions in the 3-D space [34]. High-intensity near-infrared light-emitting diodes with 850 nm wavelength (ELIMEC LTD, Israel) provided backlight to the cameras. The selected wavelength is well above the upper threshold for visual perception of most insects (∼700 nm) [35]. We did not observe any directional bias during flight of the tested animals towards or away from the light source. Animals left the glass vial one at a time through a small, inverted funnel at the opening of the vial and voluntarily started wing flapping in stagnant air. The outlet of the funnel was ∼0.35 cm (∼5 body lengths) below the cameras’ field of view to exclude transient take-off behaviours. We recorded high-speed videos from 11 animals and 36 wing stroke cycles. Due to the extreme difficulty to keep the animals within the focal range of the cameras, however, we eventually analysed only seven flapping cycles from three animals flying without visible changes in flight altitude and yaw heading. Wing and body motions were extracted from the videos using a MATLAB script published previously [36, 37]. The software derives 3-D wing position and orientation from a manually fitted 2-D computer model of the left and right contours of the membranous portion of *E. mundus* forewings (Fig. 2). The 2-D computer models assume rigid wings. Body position and orientation are calculated from four morphological markers: centre of the head, left and right wing hinges and the tip of the abdomen. From head and abdominal markers, we estimated body pitch angle between the longitudinal body axis and the horizontal (Fig. 3).

**Figure 2.**
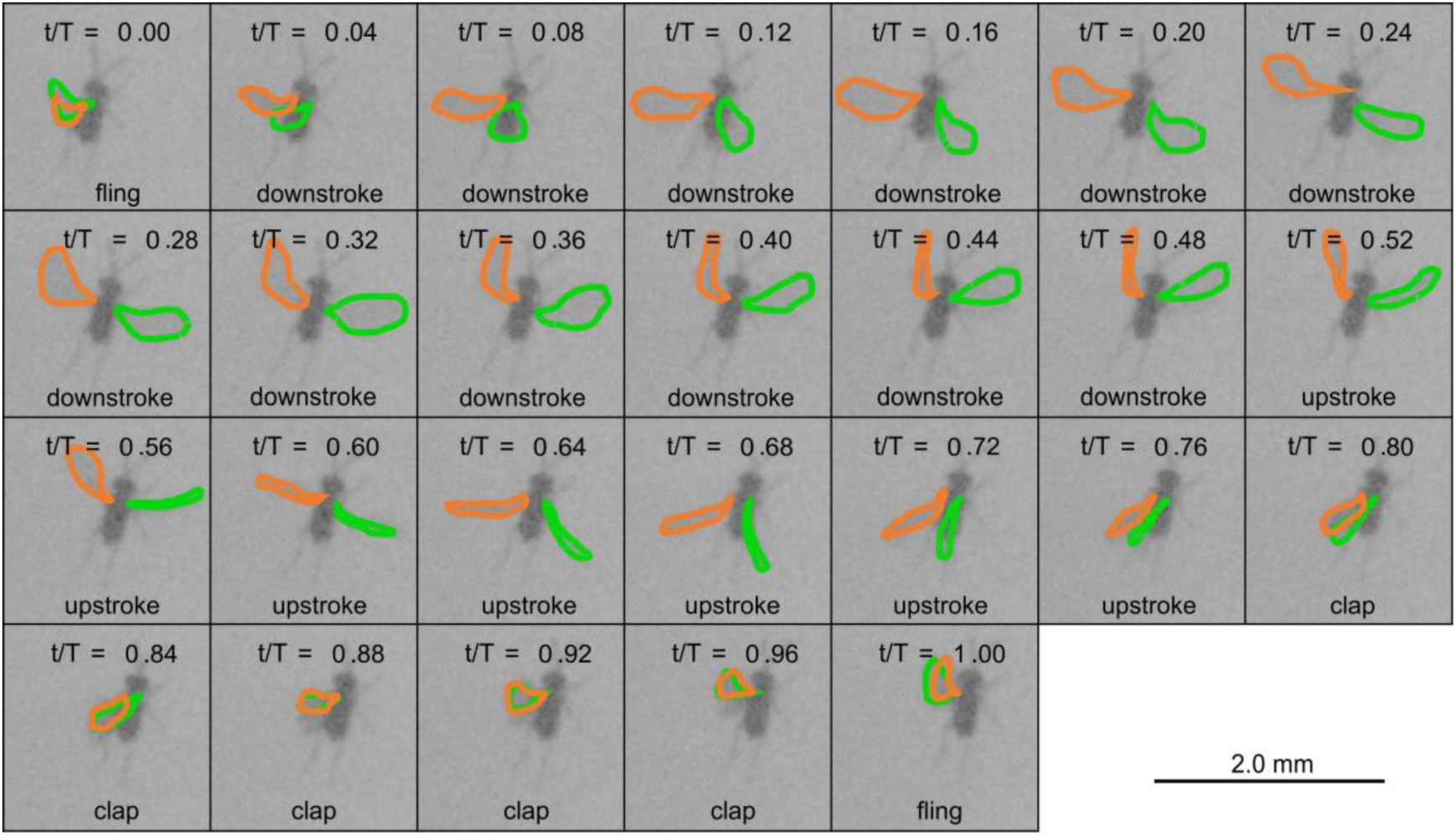
High-speed video images of a single stroke cycle in a freely flying wasp. Sequence shows 26 images recorded at 6,000 frames per seconds (∼0.167 ms per frame). Wing contours of left (orange) and right (green) forewings (membranous area) are superimposed. The stroke cycle starts with the fling that marks the beginning of the downstroke. t/T, non-dimensional time within the flapping cycle (0 - 1).

**Figure 3.**
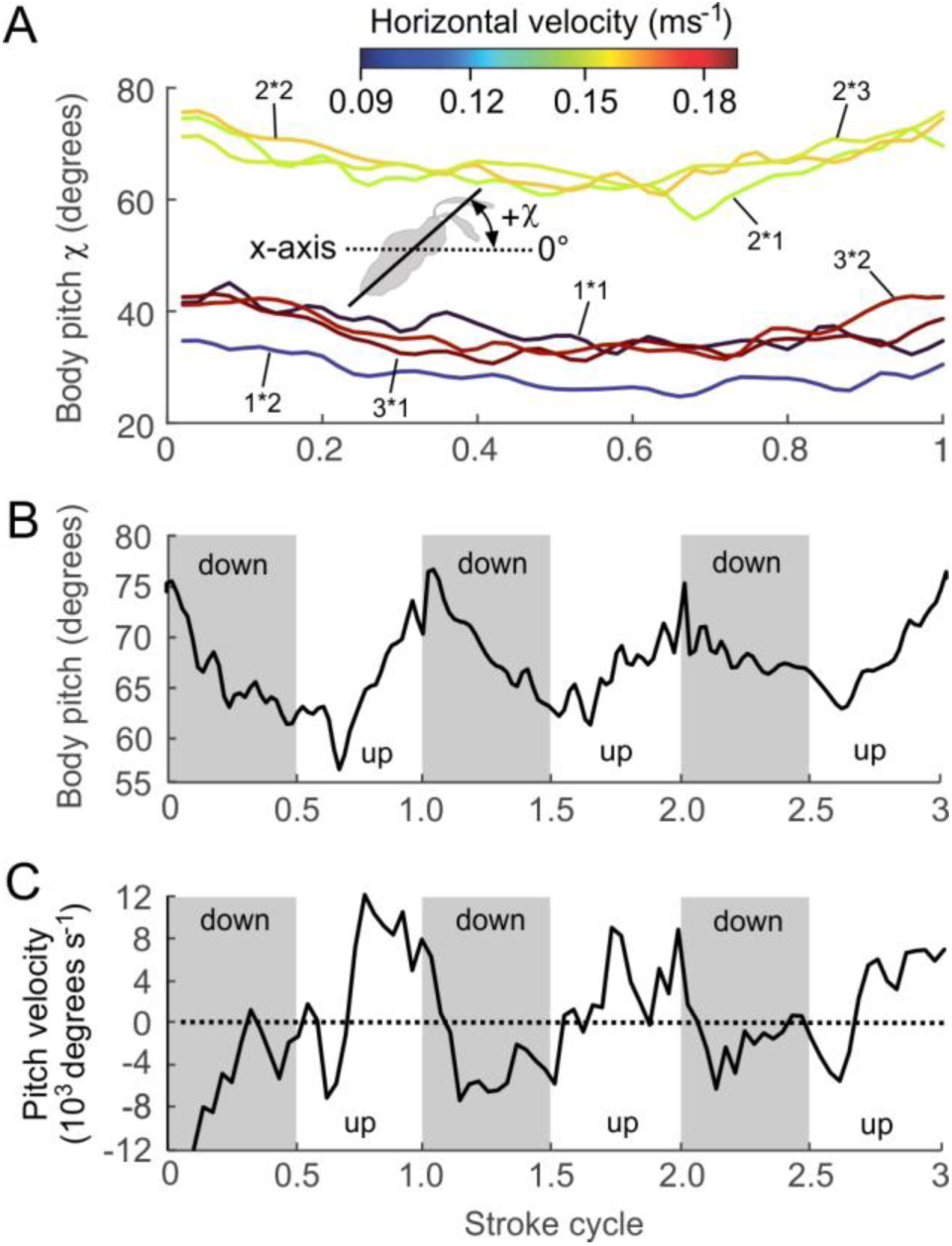
Changes in body pitch with changing flight velocity. (*A*) Oscillations of body pitch angle within a single stroke cycle. Flight sequences (cf. Table 1) are colour-coded according to the animal’s mean horizontal flight velocity (n=7 flight sequences, N=3 animals). t/T, non-dimensional time. (*B*) Changes in body pitch angle (n=3 stroke cycles and (*C*) alterations in body pitch velocity within the same 3 consecutive stroke cycles. For data smoothing see materials and methods section. Grey area indicates the duration of the downstroke.

### 2.3. Data analysis

We interpolated the video-tracked data to 50 equally-spaced time steps (2% step width) for each wingbeat cycle using the cubic-spline function in MATLAB. Position data for calculation of instantaneous body and wing velocities were smoothed by an 11-point moving average and velocities mathematically derived according to a previously published procedure [38]. Throughout the manuscript, we distinguish between three types of data representation to highlight different aspects of propulsion. (1) Figure 4 shows wing motion measured relative to a horizontally aligned body in a body frame of reference (non-inertial). (2) Figures 3 and 5 show body and wing motion in an external frame of reference (horizontal xy-axes, vertical z-axis, inertial). (3) For CFD simulation in figures 6 - 8, we use wing motion in an inertial frame of reference and relative to the insect body. In the figures, wing kinematics is presented as the average of left and right wing motion but CFD analysis used wing motion from each body side.

**Figure 4.**
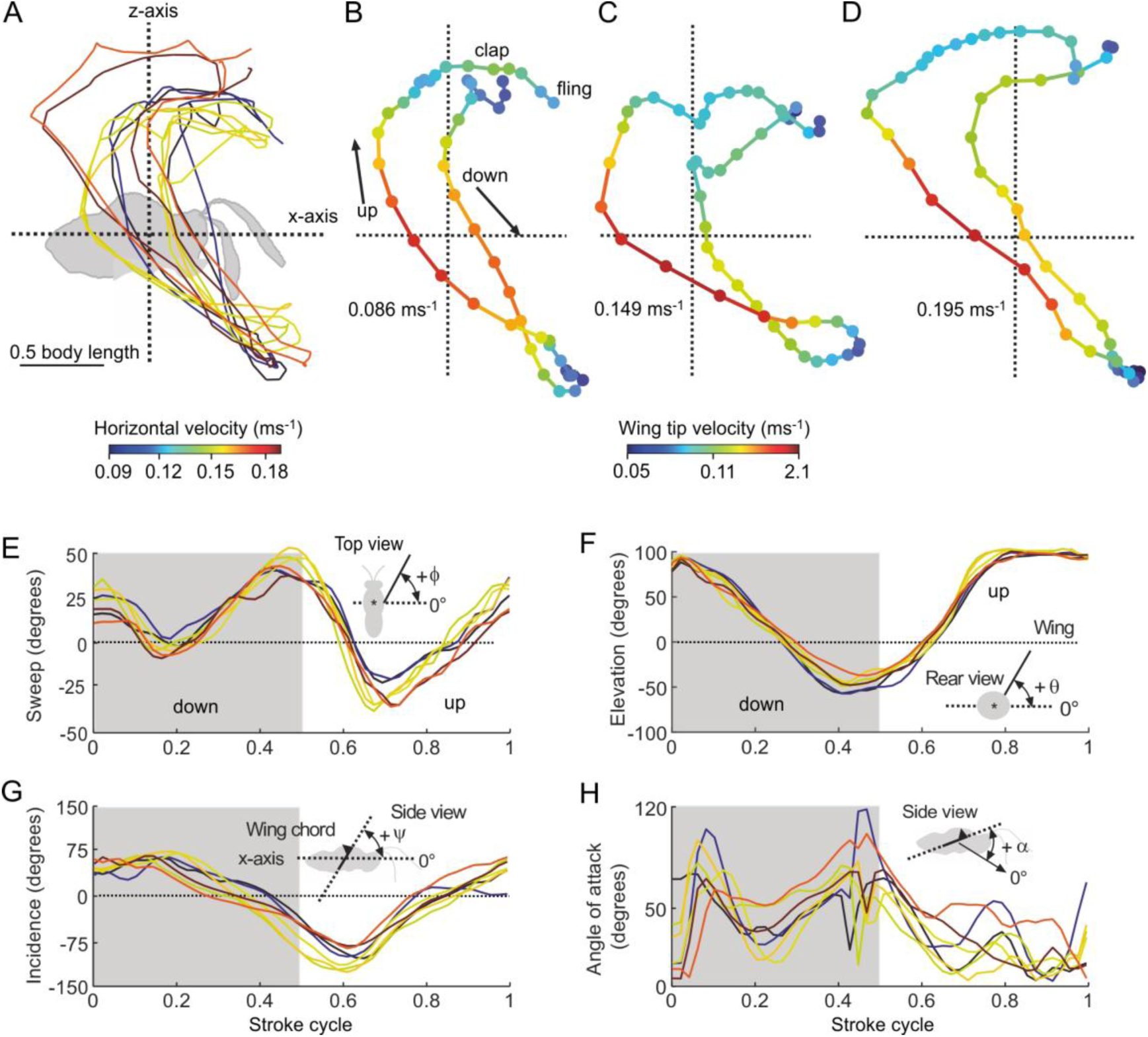
Wingtip trajectory and wingbeat parameter in the body frame of reference. (*A*) Wing tip trajectories in the sagittal plane of all analysed flapping cycles (cf. Table 1). Trajectories are means of left and right forewing with the wing hinge at origin of the coordinate system. (*B-D*) Wingtip trajectories at low (0.086 ms^-1^, 1*1), middle (0.149 ms^-1^, 2*3) and high (0.195 ms^-1^, 3*1 animal*stroke cycle) cycle-averaged horizontal forward velocities. (*E*) Wing sweep angle, (*F*) wing elevation angle, (*G*) wing incidence with respect to the horizontal, and (*H*) wing morphological angle of attack with respect to the direction of wing motion. Mean horizontal flight velocity is encoded in colour. N=3 animals, n=7 flight sequences.

**Figure 5.**
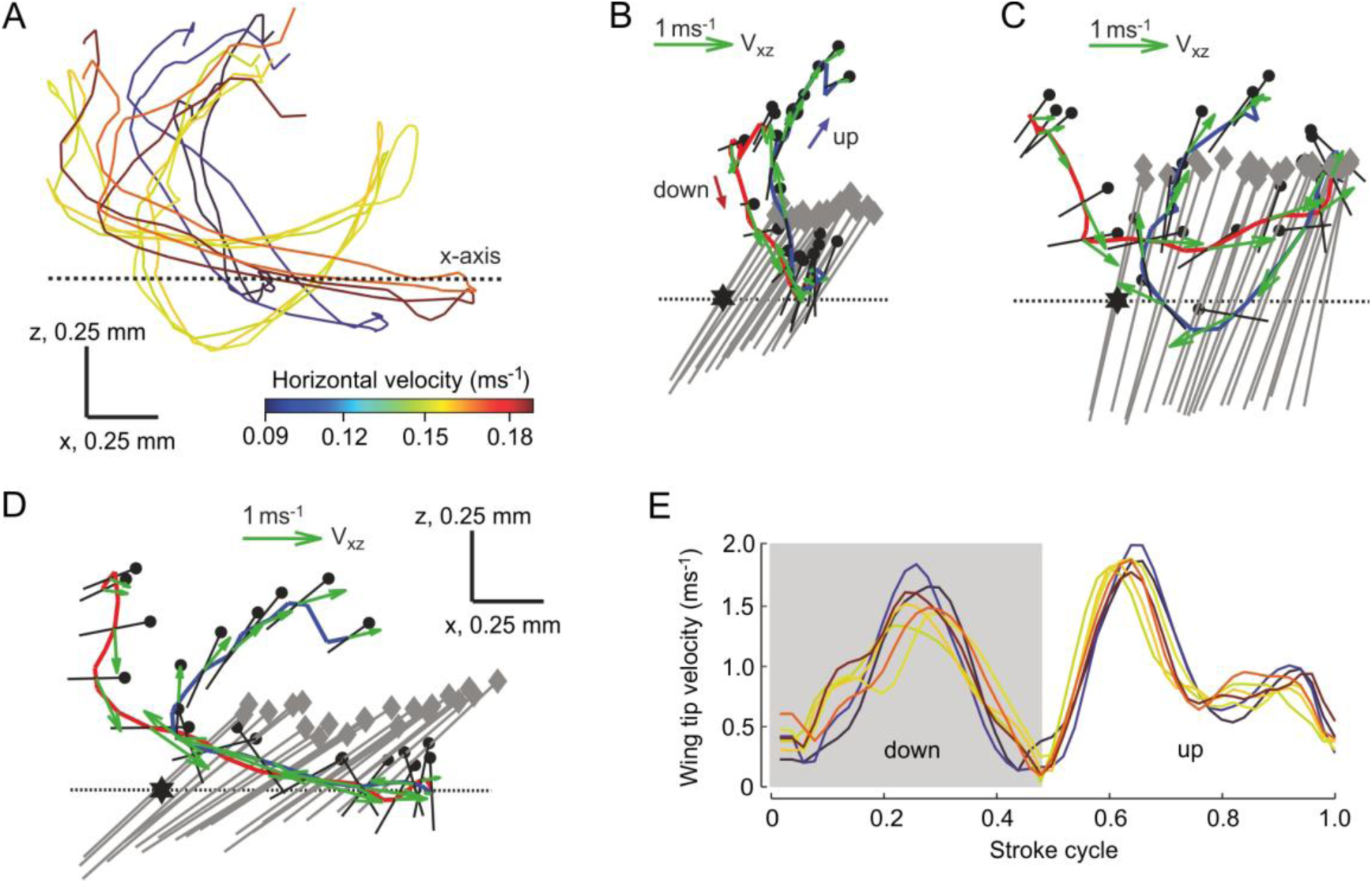
Wingtip trajectory and wing velocity in the external frame of reference. (*A*) The wing tip trajectories in the sagittal plane while the animal flies forward and upward (cf. Table 1). Trajectories are means of left and right forewing. (*B-D*) Data replotted from figure 4*B-D* but including body motion. Forward velocity in *B* is 0.086 ms^-1^, 1*1; in *C* 0.149 ms^-1^, 2*3; and in *D* 0.195 ms^-1^, 3*1 animal*stroke cycle. Wing chord is visualized at 0.6 forewing length by a black line, the wing’s leading edge as a black dot, the longitudinal body axis as grey line and the wasp’s head as a grey diamond. Green arrows indicate instantaneous wing tip velocity. Red, downstroke path; blue, upstroke path; *, position of wing hinge in the first image of the sequence. (*E*) Wingtip velocity of all flight sequences. Colours encode mean horizontal velocity of the body in *A* and *E*.

**Figure 6.**
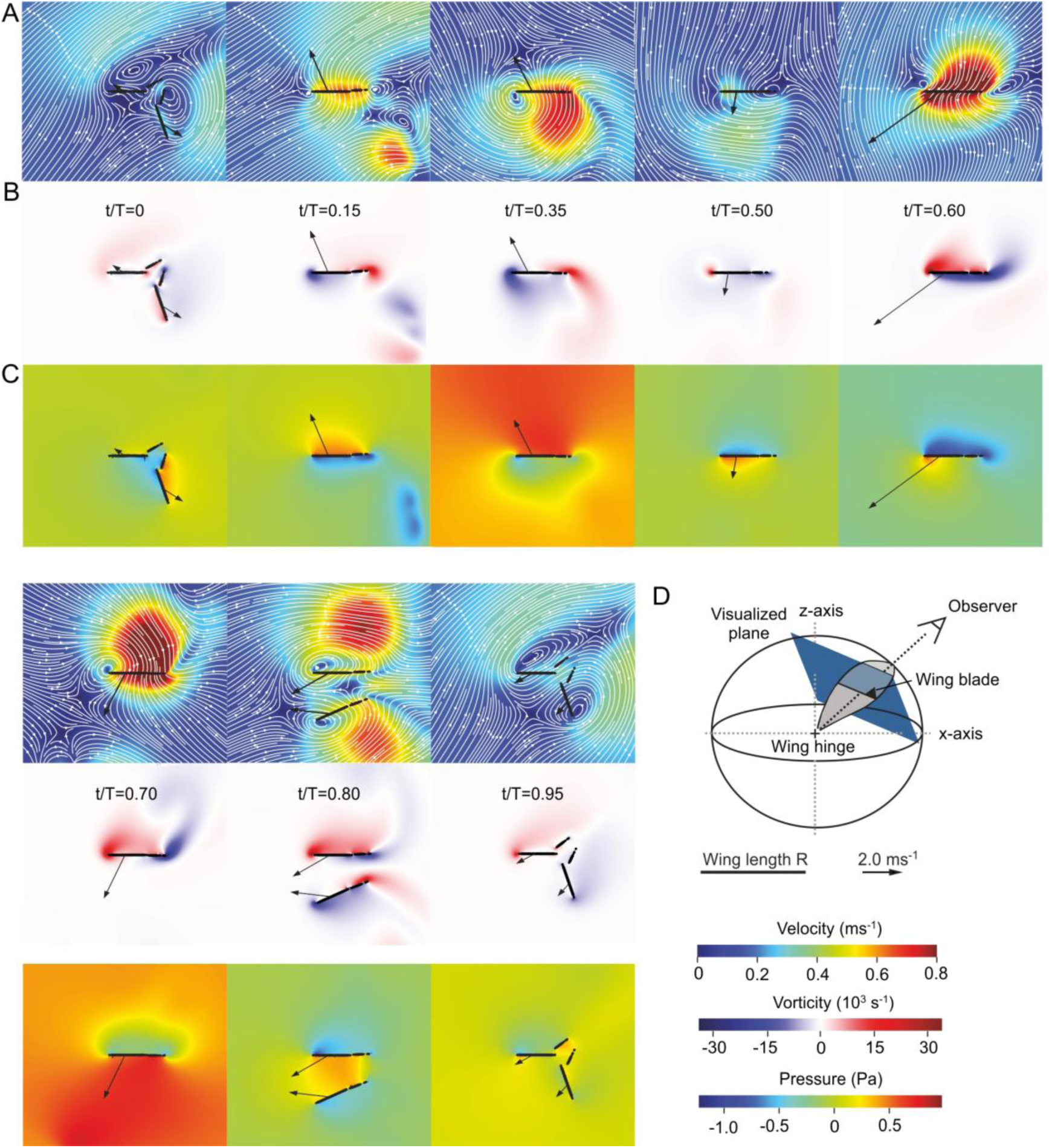
Flow visualization generated by computational fluid dynamics (CFD). Maps depict ipsilateral 2D slices from 3D data normal to the right forewing’s longitudinal axis. (*A*) Air velocity, (*B*) vorticity, and (*C*) pressure at the wing and in the wake. t/T is the normalised time within the stroke cycle, starting with the downstroke. The smaller wing chord is the hindwing and black arrows depict wing velocity. The computation shows data from animal 3 and stroke cycle 2. (*D*) Orientation of the visualized planes as shown in *A-C*. To improve visibility, the upper values of the colour bars are truncated at 50%, 25% and 35% of the maximal observed value from all animals, respectively.

All angles, equations and calculations of the three time-varying Euler angles for wing motion in the body frame of reference are described in detail in the electronic supplementary material S2. In short, wing data were rotated about each hinge into the body frame of reference and the following Euler angles derived: (1) wing elevation angle (θ) i.e. the angle between the wing’s longitudinal axis and its projection on the horizontal plane; (2) horizontal flapping angle (sweep, φ) i.e. the angle between the wing longitudinal axis and the body pitch axis after rotating the wings by θ around the body longitudinal axis; and (3) angle of incidence (ψ) i.e. the angle between wing chord and horizontal plane after rotation by θ and φ. The morphological angle-of-attack is the angle between wing chord at 0.7 wing length and direction of wing motion. To allow comparisons between stroke cycles, we normalized time (*t*) by wing cycle duration (*T*) measured from the video, starting with the beginning of the down stroke (fling motion, Fig. 2). Throughout the manuscript, data are shown as means±standard deviations.

### 2.4. Computational fluid dynamics

Our CFD simulation includes the insect body and left and right wing pairs. The numerical code 1. [39] and its validation for low Reynolds number flight were previously published [40, 41]. Here we only provide a brief summary. The CFD model is based on a finite-difference solution of incompressible Navier-Stokes equations in the artificial compressibility approximation [42, 43]. We thus numerically solve the following system of partial differential equations:

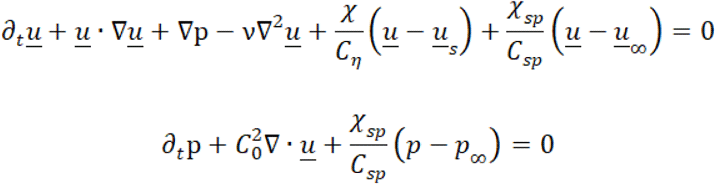

where *u̱* is the fluid velocity, p the pressure, *v* kinematic viscosity and *C_0_* artificial speed of sound. The artificial compressibility method relaxes the incompressibility constraint 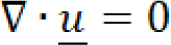 and introduces a finite speed of sound. It can, under some circumstances, be shown to be equivalent to the lattice Boltzmann method [43–45]. It is second order accurate in *C_0_* and we use a dimensionless value of *C_0_* = 20 for the simulations. The inhomogeneous Dirichlet boundary conditions are imposed using the volume penalization method [46] this method forces *u̱* towards *u̱_s_*, which is the velocity inside the solid body which is given by the rigid body equations [31]. The indicator function χ is zero in the fluid and one inside the solid (wings, body), with a smooth transition between the two [47]. The volume penalization method belongs to the class of immersed boundary methods and specifically the diffuse interface methods [48]. The outflow boundary conditions are imposed with the same method (χ*_sp_*, *C_sp_*) to impose 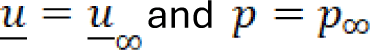 in a layer surrounding the domain, the shape of which is optimized to minimize reflections of pressure waves back into the physical domain. The two parameters *C_n_* and *C_sp_* are determined based on convergence properties [39] as a function of the minimal lattice spacing Δ*x*. This way, the volume penalization method is formally between first and second order accurate [49].

The above Navier-Stokes equations are solved with classical explicit centered finite difference of order four, combined with a Runge-Kutta-Chebychev (RKC) time stepping method, which is second order accurate [50]. Because of the combination of elevated viscosity, high local resolution to resolve individual bristles, and large domain size (we use a cube of 12-times the wasp’s wing length, 12R×12R×12R in all runs), the discretized equations are very stiff and thus challenging to solve, which may seem counter intuitive because of the low Reynolds number. The family of RKC schemes, despite being explicit in nature, are well suited for the time integration of such a stiff problem. For each simulation, we first compute the eigenvalues of the one-dimensional discrete operator in preprocessing and rescale them to three spatial dimensions. Then, using linear stability analysis, the most efficient stable RKC scheme is selected for time integration [40, 23] given the time step Δ*t* based on the Courant-Friedrichs-Lewy condition for *C*_0_. Despite being formally second order accurate, the time discretization error is negligible owing to the small time-step.

We combined the finite-differences with wavelet-based adaptivity [51] using Cohen-Daubechies-Feauveau (CDF40) wavelets to create a dynamically adaptive mesh for the spatial discretization. In every time step, the grid is adapted to the solution in order to maximize computational efficiency; it is refined where necessary and coarsened where possible. The grid is represented as octree data structure with the leaves being blocks of isotropic size of 24 grid points. This type of grid is thus a hybrid between a sparse and a dense representation as it relies on locally Cartesian blocks. Adjacent blocks may not differ in resolution by more than one level, i.e., twice or half the resolution. The CDF40 wavelet is appropriate for the high-viscosity flows studied here. The vicinity of the insect is always resolved with the maximum resolution Δ*x* allowed: for the simulation we have 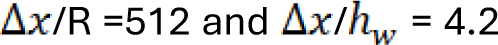, where *h_w_* is the wing thickness. The code itself has been validated in previous publications also in flow configurations relevant to the present work (i.e, including bristles), but we still performed a grid independence study (see electronic supplementary material S2, Fig S4) to ensure that results are independent of Δ*x*.

Despite the adaptive strategy, the cost of a single simulation was still approximately 30k CPUh on 400 CPU cores. To our knowledge, our code is unique for the simulation of bristled wings flapping at low Reynolds number because our adaptive strategy allows resolving the flow around each individual bristle. Our code is open source (https://github.com/adaptive-cfd/WABBIT) and freely available. All required input data, the raw output and a copy of the code version used in this work can be found in (https://osf.io/f6ztn/). For each simulation, two flapping cycles were computed, because the third cycle was found to be virtually identical to the second one. Reynolds number for wing flapping based on mean wing chord and wing tip velocity varied between ∼19 and ∼21 (Table 1).

**Table 1.**
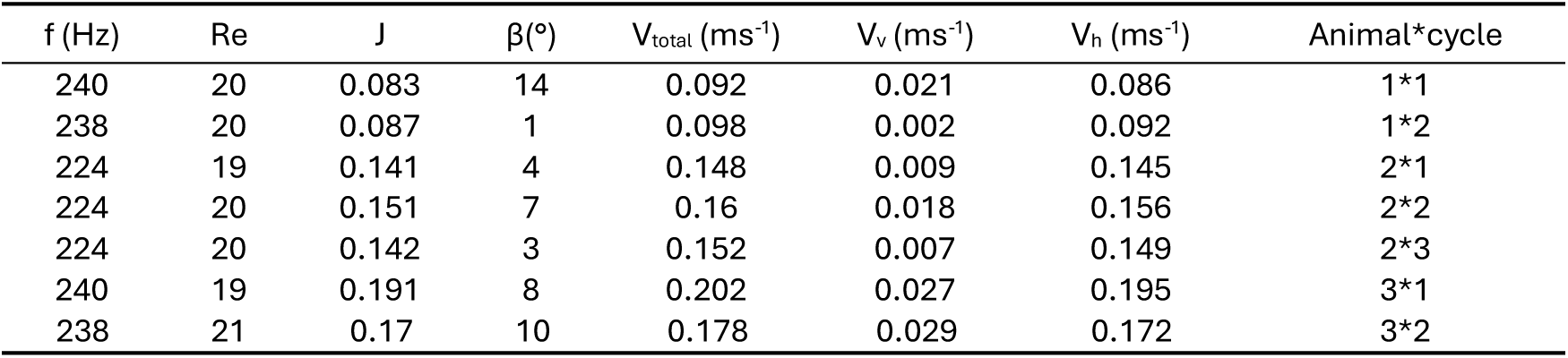
Body and wing kinematics of freely flying wasps. f, wingbeat frequency (n=5 stroke cycles including two cycles prior to and two after the analysed cycle); Re, Reynolds number calculated from mean wing tip velocity and mean wing chord; J, advance ratio; β, climbing angle i.e. the angle of the vector of flight direction with respect to the horizontal (positive angles are ascending flight); Vtotal, mean total body velocity; Vv, mean vertical body velocity; Vh, mean horizontal body velocity; animal*cycle, animal and recorded stroke cycle number.

Compared to forewings (dimensionless wing area, *A*=wing area/R^2^≈0.20, where R is forewing length), the hindwings (A≈0.038) are considerably smaller and less conspicuous in the video images, making them impossible to track (Fig. 1B, electronic supplementary material S1). We circumvented this problem by assuming that the fore-and hindwing are rigid and move as a single wing that follows forewing kinematics. The forewing kinematics, however, resulted in temporary collisions of the ipsilateral wings during fling at the dorsal stroke reversal. We avoided these collisions by rotating the hindwing about the forewing trailing edge (Fig. S1) keeping the additional rotation as small as possible. This procedure is reasonable because the wasp’s hindwings are mechanically coupled to the forewing trailing edge. We converted body and wing angles to the notation previously used for CFD analyses of flapping flight, and eventually periodized and smoothed them by restricting them to five Fourier modes. This removed the high-frequency components in the wing data owing to measurement noise [31]. The animal was simulated as a stationary object that experiences a free stream flow velocity equal to mean 3-D flight speed, i.e. a Gallilean change of reference frame. Free stream velocity is equal to negative mean body speed averaged over one complete wing stroke cycle. The animal’s body angles yaw, pitch and roll were modelled as a first order Fourier approximation, i.e. mean and one harmonic cosine and sine component fitting best the experimental data (Table S1). The simulated fluid forces were eventually decomposed into contributions from wings and body. As active motion of the insect body and legs during manoeuvring is negligible, the reported power only equals the energetic requirements to move the wings against the resisting fluid forces.

## 3. Results

### 3.1. Morphometrics and body kinematics

The wasp has membranous wings with a crown of long bristles along their margins (Fig. 1, see also electronic supplementary material S1). We found that mean forewing length (R) is ∼660±84 μm (N=7 animals) and mean length of forewing bristles ∼0.07±0.0298R. Bristle diameter at the membranous anchor point is ∼1.2±0.234 μm or ∼0.002R (n=30 bristles, N=1 wing). Mean wing thickness is ∼0.04R. Each forewing has 81 bristles and the distance between neighbouring bristle tips is ∼0.0125±0.0073R. The latter values, including bristle length, were derived from figure 8 of a previously published study [52]. We found that mean body length (L) is ∼597±82 μm (N=6 animals), body width ∼0.39±0.05L, and body height 0.25±0.03L. Mean body mass as estimated from the ellipsoid body model is ∼7.6 μg.

After taking-off from the outlet of the starting funnel, the animals reach mean horizontal and vertical velocities of ∼0.142±0.04 ms^-1^ and ∼0.016±0.01 ms^-1^, respectively (Table 1). These values convert into a velocity of ∼238 (horizontal) and ∼27 (vertical) body lengths s^-1^. Mean total flight velocity was ∼0.147±0.04 ms^-1^ and advance-ratio increases with increasing total flight velocity from ∼0.083 to ∼0.191 (Table 1). In contrast to previous studies on flying insects, mean body inclination (body pitch angle) of the wasps is not correlated with horizontal, vertical and total body velocity (Pearson correlation, -0.32<r<0.23, p>0.05, Fig. 3A, Table 1). Moreover, figure 3B shows that despite of high viscous damping due to low Reynolds number, body pitch consistently oscillates with the frequency of the flapping cycle, decreasing (increasing) pitch angle during the downstroke (upstroke). Body pitch velocity peaks at an amplitude of approximately ±12×10^3^ deg s^-1^ prior to the dorsal stroke reversals (Fig. 3C). The latter finding suggests the generation of maximum instantaneous pitching moments when wings reverse flapping direction.

### 3.2. Wing kinematics (body frame of reference)

The wing tip path relative to the insect body is U-shaped in both the upstroke and downstroke. In the body frame of reference of the horizontally aligned animal, the upstroke trajectory was consistently posterior to the downstroke at all tested flight velocities (Fig. 4). This leads to a crescent-shaped wing tip trajectory for the entire flapping cycle. Figure 4A shows the wing trajectory of all stroke cycles at various horizontal flight velocities and figure 4B-D the instantaneous wing tip velocity of three animals flying at ∼0.086, ∼0.149 and 0.195 ms^-1^ horizontal velocity, respectively. The wasps flap their wings at a mean frequency of ∼233±8 Hz (Table 1). The wing’s downstroke (∼0.04<t/T<∼0.49) was significantly longer than the upstroke (∼0.49<t/T<∼0.79), excluding clap-and-fling phase (dependent *t*-test, p<0.001). Clap-and-fling wing movements at the dorsal stroke reversal start and end on average at ∼0.79 and ∼0.04 t/T, respectively, indicating cycle-advanced wing rotation. Previous research highlighted that an advanced wing rotation maximizes force production at the stroke reversals in fruit fly-sized insects [9]. Unlike in fruit flies, however, wing rotation of our wasps pauses during the clap for ∼0.21±0.01 t/T or ∼0.9±0.05 ms. After clapping, the wings quickly separate (fling) within only ∼0.04±0.002 t/T or ∼0.16±0.006 ms.

Figure 4E-H shows the temporal changes in wing flapping angles (Euler angles) in the body reference system and the wing’s morphological angle of attack. The crescent-shaped wing trajectory leads to two maximum peaks and two minima of the wing’s sweep angle, while elevation angle and angle of incidence only peak once near the dorsal stroke reversal. None of the peak-to-peak amplitudes of elevation angle and angle of incidence are correlated with forward velocity of the animal (Pearson correlation, -0.71<r<0, p>0.07, n=7 stroke cycles, N=3 animals), but both sweep angle minima reached a lower value at faster flight speed (Pearson correlation, r<-0.87, p<0.01, n=7 stroke cycles, N=3 animals). The wing’s morphological angle of attack distinctly varies and fluctuates throughout the stroke cycle. Some of this variance is likely due to the difficulty to estimate the exact rotational angle from the video images. Mean angle peaks during the stroke reversals and range from ∼10° to 50° during the up- (0.6<t/T<0.7) and downstroke (0.2<t/T<0.3). The latter value is close to angles expected for maximum lift production in a root translating (revolving) wing (∼45°).

### 3.3. Wing kinematics (external frame of reference)

Figure 5 shows wing motion in the external frame of reference that includes motion of the body. This view is relevant for the production of aerodynamic forces and moments. Similar to figure 4, figure 5A shows the wing kinematics as a function of horizontal body velocity, and figures 5B-D display wing kinematics for three animals flying forward at ∼0.086, ∼0.149 and ∼0.195 m s^-1^, respectively. Due to the forward motion of the body, wing velocity increases during the downstroke and decreases during the early upstroke. This increasingly distorts the path trajectory compared to the trajectories shown in figure 4. Wing motion at the end of the downstroke and beginning of the upstroke is more horizontal at elevated flight velocity while flapping direction at the beginning of the downstroke and end of the upstroke remains approximately vertical. The transition from vertical to horizontal wing motion and vice versa occurs at the sweep angle minimum, at which the wing reaches the most posterior position relative to the body. At lowest flight speed, wingbeat amplitude is smallest and the wing path trajectory limited to vertical motion with pronounced differences in the wing’s angle of attack between both half strokes (Fig. 5B). We estimated a cycle-averaged wing tip velocity in the external frame of reference of ∼1.095±0.024 ms^-1^ with peak values of ∼2.0 ms^-1^ during the up- and ∼1.8 ms^-1^ during the downstroke (Fig. 5E). Despite forward body motion, wing tip velocity remains ∼25% higher in the upstroke than the downstroke yielding on average ∼1.1±0.05 ms^-1^ and ∼0.86±0.04 ms^-1^, respectively (dependent *t*-test, p<0.001, n=7 stroke cycles). Mean tip velocity in the external frame of reference is not correlated with horizontal flight velocity (Pearson correlation, r=-0.57, p=0.18), potentially suggesting a mechanism for thrust production other than wing velocity.

### 3.4. CFD simulation (flows, pressure and forces)

CFD simulation was performed for all quantified stroke cycles (Table 2). From the simulated flow data, we eventually calculated instantaneous aerodynamic forces, moments, and power. The wing blade in figure 6 is visualized at 0.6 wing length and the shown plane aligned perpendicular to the wing’s longitudinal axis (Fig. 6D). The data show the temporal evolution of air velocity (Fig. 6A), vorticity (Fig. 6B) and pressure (Fig. 6C) at the wings throughout a single stroke cycle (2*3, Table 1) and in the right forewing frame of reference. Left/right and fore-/hindwings are visible as four separate chords during the clap-and-fling manoeuvre (t/T=0, t/T=0.8-0.95). The shorter blade is the hindwing at these instants. Wing velocity vectors are indicated as black arrows. The data show that peak velocities of the flow reach up to ∼1.6 ms^-1^ and occur shortly after the ventral stroke reversals at 0.5<t/T<0.7. At this moment, the wing moves backwards (Fig. 5A,C,D) at a large ∼90° angle of attack (Fig. 4H). During the downstroke, the wings typically develop pronounced counter-clockwise vorticity at the leading wing edge (blue, t/T=0.35) and clockwise trailing edge vorticity. As expected, vorticity and flow velocity maps show that at elevated angle of attack (e.g. t/T=0.50) vorticity is concentrated near the shear layer of the leading and trailing edges, at low-pressure regions (Fig. 6C). At lower angle of attack (e.g. t/T=0.35), negative (counter clockwise) and positive (clockwise) vorticity are visible above and below the wings, respectively. The thick boundary layers suggest substantial viscous drag.

**Table 2.** Aerodynamic forces and power for flight in *E. mundus*. Pmech, mean flight muscle mass-specific mechanical power; Paero, mean aerodynamic power; Frel, ratio between total force and body weight; Ftotal, vector sum of Fx and Fz; Fz, mean vertical flight force on the body; Fx, mean horizontal force on the body; χ, body pitch angle; Mpitch, mean pitching moment; animal*cycle, animal and recorded stroke cycle number.

| $P_{mech}$ ($WKg^{-1}$) | $P_{aero}$ (nW) | $F_{rel}$ (%) | $F_{total}$ (nN) | $F_z$ (nN) | $F_x$ (nN) | $\chi$ (deg) | $M_{pitch}$ (pNm) | Animal*cycle |
| --- | --- | --- | --- | --- | --- | --- | --- | --- |
| 103.5 | 236 | 96 | 71.6 | 71 | 9 | 36.8 | 2.9 | 1*1 |
| 127.1 | 290 | 116 | 86.4 | 83 | 24 | 28.7 | -1.8 | 1*2 |
| 122.4 | 279 | 128 | 95.7 | 81 | -51 | 65.3 | 9.9 | 2*1 |
| 110.5 | 252 | 117 | 87.5 | 75 | -45 | 67.2 | -2.4 | 2*2 |
| 115.4 | 263 | 155 | 115.6 | 106 | -46 | 67.4 | -3.0 | 2*3 |
| 117.1 | 267 | 123 | 92 | 92 | 1 | 35.2 | 0.9 | 3*1 |
| 128.5 | 293 | 127 | 94.5 | 94 | 10 | 35.3 | 6.3 | 3*2 |

In figure 7A and B, we superimposed instantaneous aerodynamic forces on the processed wing kinematics in the CFD-frame of reference. The data show that the total force vector at the wing is typically tilted backwards. This reflects the contribution of viscous drag at low Re number, in turn increasing the cost for wing flapping. The total force vector was only perpendicular to the wing chord at the beginning of the downstroke (fling) and ventral stroke reversal at which the angle of attack is ∼90°. From the instantaneous forces produced by all wings and the body, we derived instantaneous (Fig. 7C, D) and mean (Table 2) horizontal thrust (F_x_) and vertical lift (F_z_). The simulation shows that thrust is scattered around zero with mean ∼-11±3.9 nN (n=7 stroke cycles). The latter suggests body acceleration when F_x_ is positive (animals 1 and 3) and breaking when F_x_ is negative (animal 2). All insects generated cycle-averaged vertical forces (∼86±12.1 nN, n=7 stroke cycles) close to or in excess of their estimated body weight (∼74.5 nN). Vertical force production by the body without wings is small, amounting to ∼1.5±0.24 nN or only ∼1.7% of mean vertical force produced by the flapping wings (Table 2). We found that the inclination angle of the combined horizontal and vertical force vector is correlated with body pitch angle (Pearson correlation, r=0.98, p<0.001). Figure 7F suggests that zero thrust is produced at a body pitch angle of ∼40° (linear regression, y = 1.12x+44.5, n=7 stroke cycles, Fig. 7F), which is close to the ∼48° mean body pitch measured in the videos for slow forward flight (Table 2). Similar to thrust, cycle-averaged body pitching moments are scattered around zero ranging from ∼-3 pNm to ∼10 pNm (Table 2).

**Figure 7.**
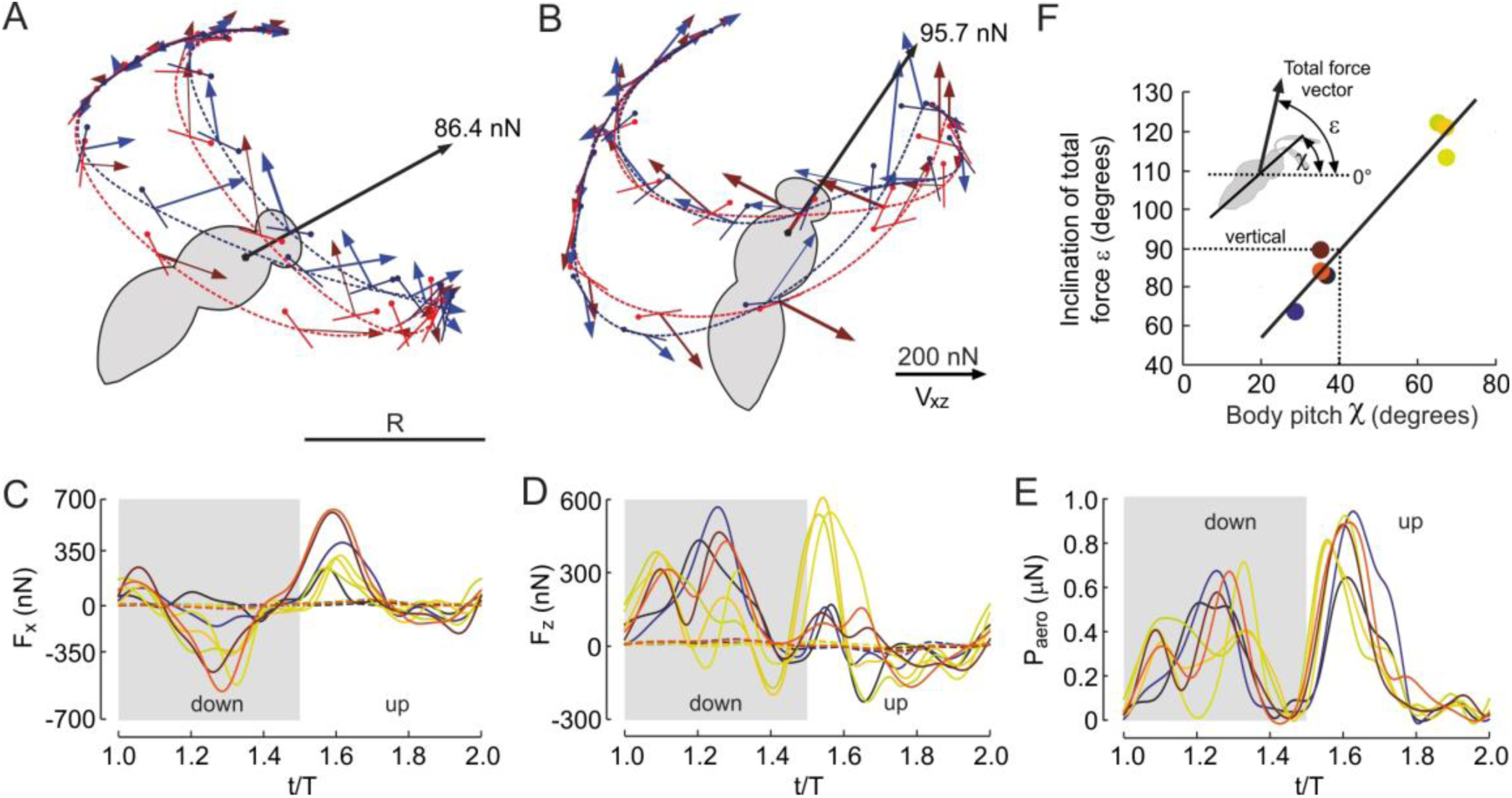
Body posture, total flight force vector, and time evolution of forces in animals flying at different horizontal velocities. (*A*) Instantaneous forces and cycle-averaged total force vector (black) and body posture of an animal flying with 0.092ms^-1^ (1*2, Table 1) and (*B*) 0.145 ms^-1^ (2*1, Table 1). Blue and red denote right and left wing, respectively. Wing positions are shown every ∼0.167 ms. (*C*) Time evolution of horizontal force (F_x_) and (*D*) vertical force component (F_z_) within one stroke cycle. Forces produced by wings are shown as solid lines and forces resulting from body drag as dashed lines. Positive values in *C* indicate forward thrust and in *d* upward force. (*E*) Time evolution of aerodynamic power within the stroke cycle assuming constant forward velocity of the body (cf. materials and methods section). Grey areas in *C*-*E* indicate the wings’ downstroke. (*F*) Inclination of cycle-averaged total force vector from each flight sequence (Table 1) determined by CFD (y-axis) with respect to the horizontal plotted against measured body pitch angle (x-axis, linear regression line, y= 1.12x + 44.5, n=7). For colour coding of mean horizontal flight velocity in *C*-*F* see figures 3 and 5*A*.

Most of the aerodynamic force in flight of *Eretmocerus mundus* is produced by the large forewings. Figure 8A and D shows that the much smaller hindwings contribute only ∼10%±1.3% and 4%±15.5% (n=7 stroke cycles) to mean vertical and horizontal forces, respectively. Both, lift and drag components normal and opposite to the direction of wing motion, respectively, contribute to vertical and horizontal forces on the insect body. Approximately 80%±13.7% of vertical force is produced by lift (Fig. 8B), while horizontal force (Fig. 8E) predominantly depends on drag when the animal produces negative thrust (∼-47 nN, animal 2, Table 2) but solely on lift at positive thrust production (∼11 nN, animals 1 and 3, Table 2). These findings suggest that even in a viscous dominated fluid regime, lift production remains the predominant component for body weight support while lift and drag share the control of thrust in the flying miniature insect. Noteworthy, the latter finding runs counter to the idea of net forward thrust production by drag-based wing rowing as previously suggested for wing motion at Re=32 - 64 [18].

**Figure 8.**
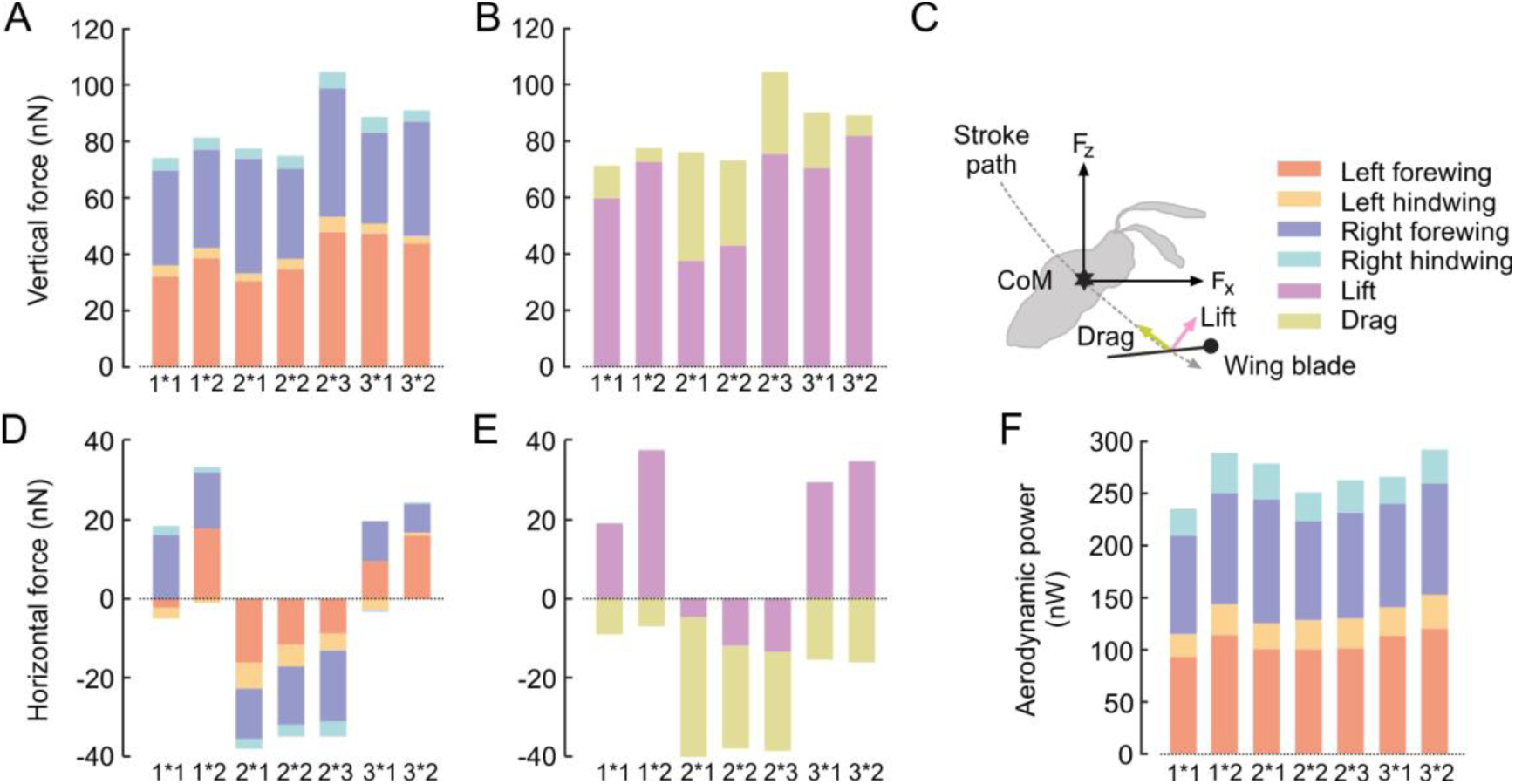
Contribution of cycle-averaged wing lift and drag to flight of *E. mundus*. (*A, D, F*) Contribution of total flight force production by each wing to vertical force (body lift, F_z_ positive value is upward) in *A*, to horizontal force (body thrust, F_x_, positive value is forward) in *D* and to aerodynamic power requirements for flight in *F*. (*B*, *E*) Contribution of lift and drag produced by all four wings to vertical force in *B* and to horizontal force in *E*. (*C*) Colour coding of data and direction of forces. X-axis, animal*stroke cycle, see Tables 1 and 2 for help; CoM, centre of mass.

### 3.5. CFD simulation (aerodynamic and muscle mechanical power)

Instantaneous wing forces computed for each flapping cycle surpass drag of the moving body by approximately two orders of magnitude (see section above). Thus, aerodynamic power requirements for flapping the wings greatly surpassed the minimal power needed for moving the body forward (Fig. 7E). Power peaks at two instants of the stroke cycle: the first peak occurs during the middle of the downstroke at t/T=∼0.3 when large aerodynamic forces are directed toward the dorsal side of the body, i.e. upward and backward relative to flight direction. The second power peak occurs shortly after the beginning of the upstroke (0.6<t/T<0.7) when total aerodynamic force points anterior, i.e. upward and forward. The peaks of aerodynamic force and power coincide with maximum wingtip velocity. In total, mean aerodynamic power among all seven stroke cycles and considering all four wings was ∼269±20.6 nW (Fig. 8F, Table 2). Assuming a 0.3 ratio between flight muscle and body mass [53], this value converts into an elevated flight muscle mass-specific mechanical power output of ∼117.8±9.01 Wkg^-1^ in *E. mundus* (Table 2). Minimum and maximum muscle mechanical power was ∼104 Wkg^-1^ and ∼129 Wkg^-1^ at 0.09 ms^-1^ and 0.18 ms^-1^ forward velocity (animals 1 and 3), respectively. As wing mass is thought to be small in miniature insects, inertial power requirements are expected to be negligible and no elastic storage in the thorax and muscle system is required [54]. We verified this assumption in more detail in the Discussion section and thus fully attribute flight muscle mechanical power output to aerodynamic power requirements.

## 4. Discussion and conclusion

### 4.1. General remarks

This study approaches the aerodynamic mechanisms and energetic costs in the tiny wasp *E. mundus* flying at low Reynolds number of ∼20. In general, it is challenging to record wing kinematics in a freely flying insect with a body length of only ∼597μm and to perform advanced computational fluid dynamics (CFD) on bristled wings because of viscous flows and computational load. The combination of both methods, however, is beneficial as it allows us to tackle kinematic behaviour and flight costs in one of the worldwide smallest flying insects at a fairly detailed level.

### 4.2. Wing kinematics and CFD modelling

The crescent-shaped wing tip trajectory in *E. mundus* has been previously reported in miniature insects such as thrips, wasps and flies [10, 22, 24, 55]. Thus, our data on *E. mundus* widely confirm the patterns of wing motion described in previous studies published on tiny and miniature insects (Figs 7 and S3). Most remarkable is that the trajectory divides wing motion into 4 phases (Fig 4B) instead of the 2 phases (up- and downstroke) typically shown for lift-based propulsion systems [9]. Our CFD modelling shows how horizontal and vertical force production is distributed within these phases (Fig. 7C and D). The main characteristics of the phases are as follows: (1) The first half of the downstroke is similar to kinematics previously described in tiny insects [10, 24] and involves elevated vertical forces due to the wing’s downward motion at high angle of attack. (2) The second half of the downstroke is similar to a conventional lift-based system, producing maximum lift at the cost of negative thrust. (3) During the first half of the upstroke, the animal predominately produces forward thrust by moving the wings backward at high angle of attack. This drag-based component is similar to wing paddling at high angle of attack and was previously described in tiny insects [55]. However, when averaged over the entire flapping cycle of *E. mundus*, drag at the wing is not responsible for forward thrust (Fig. 8E). (4) The second half of the upstroke is a vertical recovery stroke with small angles of attack and small negative lift and thrust production (Figs 5B-D, 7A-D). Wing paddling and recovery strokes have been previously shown in flight of tiny insects, including the tiny beetle *Paratuposa placentis* with 395 μm body length [10, 23, 24, 54]. Wing paddling even exist in lift-based stroke kinematics of larger insects such as *Drosophila* and may generate net forward thrust within the entire flapping cycle [28]. Despite differences in force generation mechanisms between the four wing trajectory phases, the CFD analysis shows that the outcome are distinct upstroke and downstroke peaks of force and power (Fig 7C-E), similar to the ones expected in larger insects such as *Drosophila* [36].

Altogether, miniature insects share several characteristics of the stroke cycle with their larger relatives. Many of them including *E. mundus* also enhance flight forces due to clap-and-fling kinematics [13–20, 56]. Simulation of this kinematic manoeuvre is still challenging with CFD because of several reasons. For example, clap and fling requires wing deformation that is difficult to simulate without knowing the exact biomechanic properties of the wing [57–62]. The remarkable difference in flow and vortex control between solid and deforming wings has recently been shown in a study on the fate of vortices during the dorsal stroke reversal in *Drosophila* [20]. The study shows that full clap conditions with deformable wings may alter the recycling of leading edge vortices shed at the end of the upstroke. Even though our CFD modelling would have allowed wing deformation, this approach requires the implementation of fluid-structure-interaction (FSI) and thus the biomechanical properties of the wing. Although recent studies have investigated the local mechanical properties of insect wings, no detailed data exist for wings of miniature insects [57, 63–69]. More recent progress on this problem has been made in a simulation on blowfly (*Calliphora*) wings that were modelled as a complex mass-spring network according to measurements of local wing stiffness in the fly [70]. Noteworthy, in *E. mundus*, wing bristles and thus the wing’s membranous sections are not supposed to significantly deform (up to ∼3° root-tip bristle deflection) during the up- and downstroke as suggested by a numerical study on bristles of *Paratuposa placentis* [40]. In sum, as the CFD code in this study did not permit the wings to physically touch, our estimates for maximum aerodynamic forces and power should probably be considered as a conservative approximation, excluding the contribution of full wing clapping and fling/peel motion to forces and power.

### 4.3. Body posture controls lift-thrust ratio

Large insects typically control their horizontal flight velocity by adjusting body pitch rather than changing stroke plane angle relative to the body [25]. A decrease in body pitch angle redirects the stroke plane into a more vertical orientation and the resulting total flight force vector tilts forward. Body pitching for flight speed control was previously reported for the miniature wasp *Encarsia formosa* when changing from hovering to forward flight [24] and also found in this study on *E. mundus*. However, the mechanism of force vector redirection in *E. mundus* differs from the one of larger insects. In a lift-based propulsion system with horizontally beating wings, body pitch control is largely due to vertical forces that peak owing to rotational circulation and the wake capture effect [9]. The latter forces are most effective at the stroke reversals because of an advantageous length of the moment arm. In *E. mundus*, small angles of attack during wing clapping and the elevated viscosity of air attenuate the production of vertical forces at the end of the upstroke. Walker [71] even suggested that this low force production can be viewed as a type of recovery phase in miniature insects. Figure 7A and B shows that due to the crescent-shaped wing tip path in *E. mundus*, elevated drag at the beginning of both half strokes is likely to be responsible for pitch control rather than forces at the end of each halfstroke. In this respect, the control of body pitch in the tiny wasp is delayed within the stroke cycle compared to larger insects with horizontal stroke kinematics and lift-based pitch control [9]. The two rowing phases at the beginning of each halfstroke have a counter effect on the direction of pitch rotation, though, and need to be balanced in a complete stroke cycle. As the rowing occurs twice in the stroke cycle, body pitch angle distinctly oscillates within a single stroke cycle as shown in figure 3.

### 4.4. Muscle mechanical power output

The low Reynolds number associated with body miniaturization leads to elevated energetic costs for wing flapping in insects (Fig S5, electronic supplemental materials) due to an increase in viscous drag [72]. Thus, it has previously been assumed that miniature insects require higher muscle mass-specific mechanical power output for flight than larger species. In *E. mundus* elevated viscous drag results in low instantaneous lift-to-drag ratios typically below unity for most of the stroke cycle time (Fig S6, electronic supplemental materials). The price of viscosity is the increase in aerodynamic power of up to ∼38.6 Wkg^-1^ body mass at moderate forward and climbing flight (Tables 1 and 2). Maximum locomotor capacity in *E. mundus* is thought to be higher, though, because the wasps in our experiments only reached ∼39% of their maximum flight velocity [73]. Without an increase in muscle efficiency, the increase in aerodynamic power requirements in miniature insects may lower their maximum load-lifting capacity and limit the forces available for manoeuvring [17, 74–76]. By comparison, body mass-specific aerodynamic power in large animals such as birds (e.g. starling, pigeons, ravens and gulls) and bats in cruising flight typically vary between 15 and 30 Wkg^-1^ [77–79], assuming a mechano-chemical conversion efficiency of ∼25% [80]. Body mass-specific power for flight in insects range from 20.4 to 43.2 Wkg^-1^. The smallest species tested so far are *Dasyhelea flaviventris* (0.09 mg, 29.9 Wkg^-1^), *Anbremia* sp. (0.05 mg, 20.4 Wkg^-1^), *Frankliniella occidentalis* (0.02 mg, 28.0 Wkg^-1^) and *Encarsia formosa* (0.02 mg, 25.6 Wkg^-1^) [81]. This means that flight in the ∼threefold smaller *E. mundus* is on average ∼1.4-times more costly than in *Encarsia formosa*. The above values convert into ∼68.0 to ∼100 Wkg^-1^ flight muscle mass-specific power for insects with ∼20 to ∼90 μg body mass (conversion factor=0.3, see Results section) and to ∼118 Wkg^-1^ in *E. mundus*, which is at the upper boundary of the range reported for flying animals.

Our estimates of total power requirements for flight in *E. mundus* depend on both aerodynamic and inertial power for wing flapping (Table S2, Fig. S5, electronic supplemental materials). It is quite difficult to experimentally determine wing mass in insects in the nanogram range and thus the mass is often derived from the wing’s morphology and cuticle density. For example, an elaborate study on the largely bristled winged tiny beetle *Paratuposa placentis* estimated a wing mass of ∼0.024 µg, which is approximately 1% of the animal’s body mass [23, 54]. For comparison, the latter authors suggested that a fully membranous wing with the same outline has ∼5.4 to ∼7.9 times more mass (∼0.13 μg - ∼0.19 μg) depending on its thickness [23]. The mass of membranous wings in other tiny insects is thought to be higher, amounting to ∼0.73 μg in *Trichogramma telengai*, ∼0.85 μg in the beetle *Orthoperus atomus* and ∼1.12 μg in the beetle *Limnebius atomus* [23]. For the partially membranous wings of *E. mundus*, we assumed constant wing thickness and that the membranous mass for one wing is 1% of the body mass. We then added the mass of the bristles using wing length-normalized bristle mass proposed for *P. placentis* wings (∼0.97 μg m^-1^) [23] and calculated the appropriate wing inertia tensor components (electronic supplementary material S2).

The stroke cycle-averaged inertial power in *E. mundus* is zero due to the periodic motion (the small differences shown in Table S2 are due to numerical rounding). However, the ultimate significance of inertia for total flight costs depends on the instantaneous ratio between inertial and aerodynamic power requirements and the ability of the muscle-skeletal flight system to elastically store kinetic energy [80]. Figure S4 (electronic supplemental materials) shows that in *E. mundus* instantaneous inertial power (blue) is smaller than aerodynamic power (orange) during most of stroke cycle time. At the beginning of the upstroke, inertial power peaks at 468±72 nW (N=7 cycles) that is ∼55±8.0% of aerodynamic peak power. Figure S4 further shows that there is little time within the stroke cycle of negative work during which the wasp may benefit from elastic energy storage. Thus, total power requirements without elastic energy storage are only ∼3.5% (∼8.7 nW) higher than with elastic storage (Table S2, 276 vs. 267 nW, electronic supplemental materials). In other words: *E. mundus* does not necessarily require much elastic energy storage of the flight motor to minimize flight costs because total power requirements remain positive throughout the stroke cycle [23].

Nevertheless, both inertial and aerodynamic power significantly contribute to the large peaks of total power at the stroke reversals. At the beginning of the upstroke, for example, maximum total power (1207±105 nW) exceeds cycle-averaged total power (268±21 nW, N=7, Table S2) approximately 4.5-fold in *E. mundus*, which is ∼15% more than previously reported for *P. placentis* (∼3.9) [23]. Assuming that total flight muscle mass is linked to maximum instantaneous power rather than to mean power within a stroke cycle, the elevated costs to cope with viscous drag might have caused a selective evolutionary pressure on miniature insects to minimize inertial costs and thus wing mass. The reduction in inertial power can be achieved in different ways, for example, by changes to kinematics, wing shape, a reduction in wing thickness, and by replacing some of the solid wing area by bristles [23, 41, 54, 83]. The aerodynamic performance loss in the latter case is relatively small at low Reynolds number because the thick boundary layer between bristles hinders flow to pass through the porous membrane [3, 12, 83].

### 4.5. Conclusion

Insect flight in a viscosity-dominated fluid realm differs from insect flight at elevated Reynolds numbers in several respects, such as the crescent-shaped path of wing motion and the elevated flight costs. Our CFD analysis shows how energetic costs and locomotor propulsion mechanisms are linked and associated with the extreme miniaturization of the insect body. Most of our data confirm previous results on flight in miniature insects but we also report novel details on body posture and wing control. This includes a modified clap-and-fling manoeuvre and alterations of body pitch for flight speed control by two counter-acting rowing phases at the beginning of each halfstroke. CFD-based estimates of aerodynamics power are more precise than estimates based on simple cycle-averaged kinematics. Besides the contribution of fluid viscosity, this might be one reason why our power estimates in *E. mundus* are at the upper limit of values reported so far. It is reasonable to assume that the elevated flight costs represent an evolutionary pressure on the development of bristled wings because the replacement of a solid wing membrane by bristles may reduce wing mass up to ∼8-times. Such reduction in wing mass need not come at the expense of reduced aerodynamic efficiency allowing the presence of both membranous and bristle wings in miniature insects, whereas only membranous wings allow efficient force production in larger insects [40]. Altogether, a close comparison of wingbeat control and flight costs between small and large insects should help to fully understand the entire scope of flight mechanisms in animals and thus the limits of miniaturization during the evolution of flight.

## Supporting information

Electronic supplementary material

## Data accessibility

All data needed to evaluate the conclusions in the paper are present in the paper itself and in the electronic supplementary material. The CFD solver and relevant data are available at https://osf.io/f6ztn/

## Author contributions

Conceptualization: A.S., G.R.; Methodology: A.S., T.E., G.R., F.-O.L.; Software: A.S., T.E.; Formal analysis: A.S., T.E., G.R.; Investigation: A.S.; Resources: T.E., G.R.; Writing - original draft: A.S., G.R.; Writing - review & editing: F.-O.L.; Visualization: A.S., T.E., F.-O.L.; Supervision: G.R.; Project administration: G.R., F.-O.L.; Funding acquisition: T.E., G.R., F.-O.L.

## Acknowledgments

We thank R. Rotem and E. Sar-shalom for their help in insect rearing. We are grateful to F. Muijres for access and help with modifications of the tracking code.

## List of symbols

α: wing’s morphological angle of attack
β: body climbing angle with respect to the horizontal
ε: inclination of total force vector with respect to the horizontal
χ: body pitch with respect to the horizontal
φ: wing sweep angle
ψ: angle of incidence between wing chord and animal’s horizontal plane
θ: elevation angle between wing longitudinal axis and the animal’s transverse axis
*A*: Dimensionless wing area (membranous area·R^-2^)
f: wingbeat frequency
F_x_: horizontal flight force
F_z_: vertical flight force
F_total_: total flight force
F_rel_: ratio between total flight force and body weight
J: advance ratio
m: body mass
M_pitch_: body pitch moment
P_aero_: absolute aerodynamic power for wing and body motion
P_mech_: muscle mass-specific mechanical power of the flight muscle
r: pearson correlation coefficient
R: forewing length (excluding bristles)
Re: Reynolds number
t: time
T: duration of stroke cycle
V_h_: horizontal body velocity
V_v_: vertical body velocity
V_total_: total body velocity
*u̱*: flow velocity in the simulation
*u̱_s_*: the velocity inside the solid body
p: pressure
*u̱*_∞_, *p*_∞_: free-stream velocity and pressure
*v*: kinematic viscosity
*C_0_*: artificial speed of sound
*X*,*X_sp_*: indicator function of obstacle and outflow
*C_η_*,*C_sp_*: Penalization parameters (numerical permeability) for obstacle and outflow

## References

[1] Ellington, C.P. 1999 The novel aerodynamics of insect flight: applications to micro-air vehicles. J. Exp. Biol. 202, 3439–3448.

[2] Sane, S.P. 2016 Neurobiology and biomechanics of flight in miniature insects. Curr. Opin. Neurobiol. 41, 158–166.

[3] O’Callaghan, F., Sarig, A., Ribak, G. & Lehmann, F.-O. 2022 Efficiency and aerodynamic performance of bristled insect wings depending on Reynolds number in flapping flight. Fluids 7, 75.

[4] Horridge, G.A. 1956 The flight of very small insects. Nature 178, 1334–1335.

[5] Vogel, S. 1988 Life’s Devices: the physical world of animal and plants. Princeton, Princeton University Press; 367 p.

[6] Jones, S.K., Laurenza, R., Hedrick, T.L., Griffith, B.E. & Miller, L.A. 2015 Lift vs. drag based mechanisms for vertical force production in the smallest flying insects. J. Theor. Biol. 7, 105–120. (doi:10.1016/j.jtbi.2015.07.035).

[7] Ellington, C.P., Berg, C.v.d., Willmott, A.P. & Thomas, A.L.R. 1996 Leading-edge vortices in insect flight. Nature 384, 626–630.

[8] Dickinson, M.H. 1994 The effects of wing rotation on unsteady aerodynamic performance at low Reynolds numbers. J. Exp. Biol. 192, 179–206.

[9] Dickinson, M.H., Lehmann, F.-O. & Sane, S. 1999 Wing rotation and the aerodynamic basis of insect flight. Science 284, 1954–1960.

[10] Cheng, X. & Sun, M. 2018 Very small insects use novel wing flapping and drag principle to generate the weight-supporting vertical force. J. Fluid Mech. 855, 646–670.

[11] Santhanakrishnan, A., Jones, S.K., Dickson, W.B., Peek, M., Kasojo, V.T., Dickinson, M.H. & Miller, L.A. 2018 Flow structure and force generation on flapping wings at low reynolds numbers relevant to the flight of tiny insects. Fluids 3, 45.

[12] O’Callaghan, F. & Lehmann, F.-O. 2023 Flow development and leading edge vorticity in bristled insect wings. J. Comp. Physiol. A 209, 219–229.

[13] Lighthill, M.J. 1973 On the Weis-Fogh mechanism of lift generation. J. Fluid Mech. 60, 1–17.

[14] Weis-Fogh, T. 1973 Quick estimates of flight fitness in hovering animals, including novel mechanisms for lift production. J. Exp. Biol. 59, 169–230.

[15] Ellington, C.P. 1975 Non-steady-state aerodynamics of the flight of Encarsia formosa. New York, Plenum Press; 783–796 p.

[16] Maxworthy, T. 1979 Experiments on the Weis-Fogh mechanism of lift generation by insects in hovering flight Part 1. Dynamics of the ’fling’. J. Fluid Mech. 93, 47–63.

[17] Marden, J.H. 1987 Maximum lift production during takeoff in flying animals. J. Exp. Biol. 130, 235–258.

[18] Miller, L.A. & Peskin, C.S. 2005 A computational fluid dynamics of ’clap and fling’ in the smallest insects. J. Exp. Biol. 208, 195–212.

[19] Lehmann, F.-O. 2008 When wings touch wakes: understanding locomotor force control by wake–wing interference in insect wings. J. Exp. Biol. 211, 224–233.

[20] Lehmann, F.-O., Wang, H. & Engels, T. 2021 Vortex trapping recaptures energy in flying fruit flies. Sci. Reports 11, 1–7.

[21] Kasoju, V.T. & Santhanakrishnan, A. 2021 Pausing after clap reduces power required to fling wings apart at low Reynolds number. Bioinsp. Biomim. 16, 056006.

[22] Lyu, Y.Z., Zhu, H.J. & Sun, M. 2019 Flapping-mode changes and aerodynamic mechanisms in miniature insects. Phys. Rev. E 99, 012419.

[23] Farisenkov, S.E., Kolomenskiy, D., Petrov, P.N., Engels, T., Lapina, N.A., Lehmann, F.-O., Onishi, R., Liu, H. & Polilov, A.A. 2022 Novel flight style and light wings boost flight performance of tiny beetles. Nature 602, 96–100.

[24] Cheng, X. & Sun, M. 2021 Wing kinematics and aerodynamic forces in miniature insect *Encarsia formosa* in forward flight. Phys. Fluids 33.

[25] Dudley, R. & Ellington, C.P. 1990 Mechanics of forward flight in bumblebees. I. Kinematics and morphology. J. Exp. Biol. 148, 19–52.

[26] Meng, X.G. & Sun, M. 2016 Wing and body kinematics of forward flight in drone-flies. Bioinsp. Biomim. 11, 056002.

[27] Willmott, A.P. & Ellington, C.P. 1997 The mechanics of flight in the hawkmoth *Manduca sexta*. I. Kinematics of hovering and forward flight. J. Exp. Biol. 200, 2705–2722.

[28] Ristroph, L., Bergou, A.J., Guckenheimer, J., Wang, Z.J. & Cohen, I. 2011 Paddling mode of forward fligth in insects. Phys. Rev. Lett. 106, 178103. (doi:DOI: 10.1103/PhysRevLett.106.178103).

[29] Mercet, R.G. 1931 Notas sobre Aphelinidos (Hym. Chalc.). Revista Española de Entomología 7, 395.

[30] Ribak, G., Dafni, E. & Gerling, D. 2016 Whiteflies stabilize their take-off with closed wings. J. Exp. Biol. 219, 1639–1648.

[31] Engels, T., Kolomenskiy, D., Schneider, K. & Sesterhenn, J. 2016 FluSI: A novel parallel simulation tool for flapping insect flight using a Fourier method with volume penalization. SIAM J. Sci. Comp. 38, S3–S24.

[32] Urca, T. & Ribak, G. 2018 The effect of air resistance on the jump performance of a small parasitoid wasp, *Anagyrus pseudococci* (Encyrtidae). J. Exp. Biol. 221, jeb177600.

[33] Bennett-Clark, H.C. & Alder, G.M. 1979 The effect of air resistance on the jumping performance of insects. J. Exp. Biol. 82, 105–121.

[34] Hedrick, T.L. 2008 Software techniques for two- and three-dimensional kinematic measurements of biological and biomimetic systems. Bioinsp. Biomim. 3. (doi:10.1088/1748-3182/3/3/034001).

[35] Döring, T.F. & Chittka, L. 2007 Visual ecology of aphids—a critical review on the role of colours in host finding. Arthrop. Plant Interact. 1, 3–16.

[36] Fry, S.N., Sayaman, R. & Dickinson, M.H. 2005 The aerodynamics of hovering flight in *Drosophila*. J. Exp. Biol. 208, 2303–2318.

[37] Fontaine, E.I., Zabala, F., Dickinson, M.H. & Burdick, J.W. 2009 Wing and body motion during flight initiation in *Drosophila* revealed by automated visual tracking. J. Exp. Biol. 212, 1307–1323.

[38] Rayner, J.M.V. & Aldridge, H.D.J. 1985 Three-dimensional resonstruction of animal flight paths and the turning flight of micropteran bats. J. Exp. Biol. 118, 247–265.

[39] Engels, T., Schneider, K., Reiss, J. & Farge, M. 2021 A wavelet-adaptive method for multiscale simulation of turbulent flows in flying insects. Commun. Comput. Phys., 30:1118–1149

[40] Engels, T., Kolomenskiy, D. & Lehmann, F.-O. 2021 Flight efficiency is a key to diverse wing morphologies in small insects. J. R. Soc. Interface 18, 20210518.

[41] Kolomenskiy, D., Farisenkov, S., Engels, T., Lapina, N., Petrov, P., Lehmann, F.-O., Onishi, R., Liu, H. & Polilov, A.A. 2020 Aerodynamic performance of a bristled wing of a very small insect. Exp. Fluids 61, 1–13.

[42] Chorin, A.J. 1967 A numerical method for solving incompressible viscous flow problems. J. Comp. Phys. 2, 12–26.

[43] Ohwada, T. & Asinari, P. 2010 Artificial compressibility method revisited: Asymptotic numerical method for incompressible Navier–Stokes equations. J. Comp. Phys. 229, 1698– 1723.

[44] Asinari, P., Ohwada, T., Chiavazzo, E. & Di Rienzo, A. F. 2012 Link-wise artificial compressibility method. J. Comput. Phys. 231:5109–5143.

[45] Ohwada, T., Asinari, P. & Yabusaki, D. 2011 Artificial compress ibility method and lattice boltzmann method: Similarities and differences. Computers & Mathematics with Applications 61(12):3461–3474.

[46] Angot, P., Bruneau, C. & Fabrie, P. 1999 A penalization method to take into account obstacles in incompressible viscous flows. Numer. Math. 81:497– 520.

[47] Engels, T., Kolomenskiy, D., Schneider, K. & Sesterhenn, J. 2015 Numerical simulation of fluid-structure interaction with the volume penalization method. J. Comput. Phys. 281:96–115.

[48] Sotiropoulos, F. & Yang, X. 2014 Immersed boundary methods for simulating fluid–structure interaction. Prog. Aerospace Sci. 65:1–21.

[49] Engels, T., Truong, H., Farge, M., Kolomenskiy, D. & Schneider K. 2022 Computational aerodynamics of insect flight using volume penalization. Comptes rendus mécanique.

[50] Verwer, J. G., Hundsdorfer, W. H. & Sommeijer, B P. 1989 Convergence properties of the runge-kutta-chebyshev method. Numer. Math. 57:157–178.

[51] Schneider, K. & Vasilyev, O. 2010 Wavelet methods in computational fluid dynamics. Annu. Rev. Fluid Mech. 42:473–503.

[52] Barro, P.J.D., Driver, F., Naumann, I.D., Schmidt, S., Clarke, G.M. & Curran, J. 2000 Descriptions of three species of *Eretmocerus* Haldeman (Hymenoptera: Aphelinidae) parasitising *Bemisia tabaci* (Gennadius)(Hemiptera: Aleyrodidae) and *Trialeurodes vaporariorum* (Westwood)(Hemiptera: Aleyrodidae) in Australia based on morphological and molecular data. Aust. J. Entomol. 39, 259–269.

[53] Lehmann, F.-O. & Dickinson, M.H. 1997 The changes in power requirements and muscle efficiency during elevated force production in the fruit fly, *Drosophila melanogaster*. J. Exp. Biol. 200, 1133–1143.

[54] Farisenkov, S.E., Lapina, N.A., Petrov, P.N. & Polilov, A.A. 2020 Extraordinary flight performance of the smallest beetles. PNAS 117, 24643–24645.

[55] Zhao, P., Dong, Z., Jiang, Y., Liu, H., Hu, H., Zhu, Y. & Zhang, D. 2019 Evaluation of drag force of a thrip wing by using a microcantilever. J. Appl. Phys. 126.

[56] Santhanakrishnan, A., Robinson, A.K., Jones, S., Low, A.A., Gadi, S., Hedrick, T.L. & Miller, L.A. 2014 Clap and fling mechanism with interacting porous wings in tiny insect flight. J. Exp. Biol. 217, 3898–3909.

[57] Wehmann, H.-N., Heepe, L., Gorb, S.N., Engels, T. & Lehmann, F.-O. 2019 Local deformation and stiffness distribution in fly wings. Biol. Open 8, bio038299. (doi:10.1242/bio.038299).

[58] Shumway, N. & Laurence, S.J. 2019 The impact of deformation on the aerodynamics of flapping dragonfly wings. In AIAA Scitech 2019 Forum (p. 1378.

[59] Yin, D., Wei, Z., Wang, Z. & Zhou, C. 2018 Measurement of shape and deformation of insect wing. Rev. Sci. Instrum. 89, 014301. (doi:10.1063/1.5019200).

[60] Koehler, C., Liang, Z., Gaston, Z., Wan, H. & Dong, H. 2012 3D reconstruction and analysis of wing deformation in free-flying dragonflies. J. Exp. Biol. 215, 3018–3027. (doi:10.1242/jeb.069005).

[61] Lehmann, F.-O., Gorb, S., Nasir, N. & Schützner, P. 2011 Elastic deformation and energy loss of flapping fly wings. J. Exp. Biol. **l**214, 2949–2961.

[62] Du, G. & Sun, M. 2010 Effects of wing deformation on aerodynamic forces in hovering hoverflies. J. Exp. Biol. 213, 2273–2283.

[63] Combes, S.A. & Daniel, T.L. 2003 Flexural stiffness in insect wings I. Scaling and the influence of wing venation. J. Exp. Biol. 206, 2979–2987.

[64] Combes, S.A. & Daniel, T.L. 2003 Flexural stiffness in insect wings II. Spatial distribution and dynamic wing bending. J. Exp. Biol. 206, 2989–2997.

[65] Chen, Y.H., Skote, M., Zhao, Y. & Huang, W.M. 2013 Stiffness evaluation of the leading edge of the dragonfly wing via laser vibrometer. Materials Lett. 97, 166–168. (doi:10.1016/j.matlet.2013.01.110).

[66] Mengesha, T.E., Vallance, R.R. & Mittal, R. 2011 Stiffness of desiccating insect wings. Bioinsp. Biomim. 6, 014001. (doi:10.1088/1748-3182/6/1/014001).

[67] Steppan, S.J. 2000 Flexural stiffness patterns of butterfly wings (Papilionoidea). J. Res. Lepid. 35, 61–77.

[68] Engels, T., Wehmann, H.-N. & Lehmann, F.-O. 2020 Three-dimensional wing structure attenuates aerodynamic efficiency in flapping fly wings. J. R. Soc. Interface 17, 20190804. (doi:10.1098/rsif.2019.0804).

[69] Krishna, S., Cho, M., Wehmann, H.-N., Engels, T. & Lehmann, F.-O. 2020 Wing design in flies: properties and aerodynamic function. Insects 11, 466. (10.3390/insects11080466).

[70] Truong, H., Engels, T., Wehmann, H., Kolomenskiy, D., Lehmann, F.-O. & Schneider, K. 2021 An experimental data-driven mass-spring model of flexible *Calliphora* wings. Bioinsp. Biomim. 17, 026003. (10.1088/1748-3190/ac2f56).

[71] Walker, J.A. 2002 Functional morphology and virtual models: physical constraints on the design of oscillating wings, fins, legs and feet at intermediate reynolds numbers. Integ. Comp. Biol. 42, 232–242.

[72] Vogel, S. 1994 Life in Moving Fluids. 2nd ed. Princeton, Princeton University Press.

[73] Sarig, A. & Ribak, G. 2021 To what extent can the tiny parasitoid wasps, *Eretmocerus mundus*, fly upwind? J. Appl. Entomol. 145, 660–674.

[74] Lehmann, F.-O. 2001 The efficiency of aerodynamic force production in *Drosophila*. Comp. Biochem. Physiol. A 131, 77–88.

[75] Marden, J.H. 1989 Effects of load-lifting constraints on the mating system of a dance fly. Ecology 70, 496–502.

[76] Mountcastle, A.M. & Combes, S.A. 2013 Wing flexibility enhances load-lifting capacity in bumblebees. Proc. Roy. Soc. Lond. A 280, 20130531. (doi:10.1098/rspb.2013.0531).

[77] Norberg, U.M. 1996 Energetics of flight. In Avian energetics and nutritional ecology (pp. 199–249, Springer.

[78] Ellington, C.P. 1991 Limitations on animal flight performance. J. Exp. Biol. 160, 71–91.

[79] Dial, K.P., Biewener, A.A., Tobalske, B.W. & Warrick, D. 1997 Mechanical power output of bird flight. Nature 390, 67–70.

[80] Ellington, C.P. 1984 The aerodynamics of hovering insect flight. VI. Lift and power requirements. Phil. Trans. R. Soc. Lond. B 305, 145–181.

[81] Lyu, Y.Z. & Sun, M. 2021 Power requirements for the hovering flight of insects with different sizes. J. Insect Physiol. 134, 104293.

[82] Dickinson, M.H. & Lighton, J.R.B. 1995 Muscle efficiency and elastic storage in the flight motor of *Drosophila*. Science 268, 87–89.

[83] Cheer, A.Y.L. & Koehl, M.A.R. 1987 Paddles and rakes: fluid flow through bristled appendages of small organisms. J. Theor. Biol. 129, 17–39.

