## Supplementary material for "Muscle power output reflects elevated viscosity in the propulsion system of flying miniature wasps": Electronic supplementary material

### Electronic supplemental materials

##### S1 Wing model with bristles used in the flow simulations

Figure S1 shows the outline of the wing in *E. mundus* including the reconstruction of wing bristles. These data are used for CFD modelling.

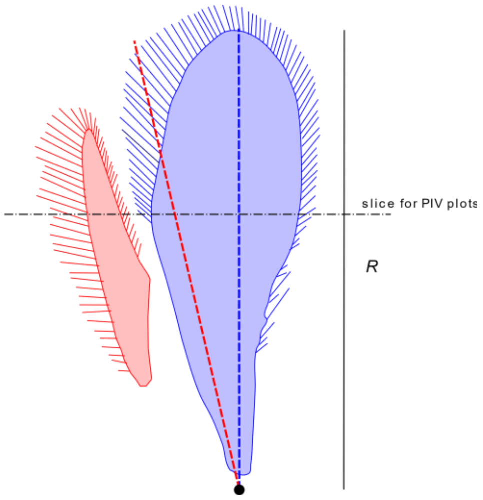


**Figure S1.** Wing morphology of the wasp. Axes are scaled by wing length (R=660 µm). Dashed blue and red lines denote the axes of wing rotation for the fore- and hindwings, respectively. Dashed black line denotes the wing section at 0.6R that is shown by CFD flow and pressure maps in figure 6.

##### S2 Extended methodology

*S2.1 Calculation of wingtip trajectory in the external frame of reference in the sagittal plane*

To compare wing tip trajectory, we only used horizontal flights. We defined the frame of reference ($X_f$,$Y_f$,$Z_f$), with the $X_f$-axis in the direction of the line connecting the body's position at the start and end of the flapping cycle. $Y_f$- and $Z_f$-axis are normal to $X_f$ with $Y_f$ being horizontal and pointing to the left of the insect. The position of the centre of the body at the beginning of the flapping cycle was subtracted from the following data points. Next, the flight direction vector (FD) was determined in the cameras' frame of reference and the coordinate system was rotated to align FD with $X_f$. All analysed flight sequences were level flights of the wasps. Rotation was in the horizontal XY-plane of the cameras and flight direction in this azimuth plane is:

$$\alpha = tan^{-1}(FD_X, FD_Y). \qquad \text{(eq. 1)}$$

Tracked data points (e.g., point $P_2$) were rotated by $\alpha$ about the Z-axis:

$$P_2{}' = \begin{bmatrix} \cos\alpha & -\sin\alpha & 0 \\ \sin\alpha & \cos\alpha & 0 \\ 0 & 0 & 1 \end{bmatrix} \times P_2. \qquad \text{(eq. 2)}$$

Data from the left and right wing of each insect were averaged and the trajectory was plotted in the vertical ($X_fZ_f$) plane of the flight coordinate system.

*S2.2 Data transformation from the camera to the body frame of reference*

The transformation of the data from the camera frame of reference to the body frame of reference was as follows:

1) In each video frame, the centre of the body in the camera frame of reference is used as the origin for the body frame of reference and the subsequent data are shifted accordingly.
2) Using the points on head, tip of abdomen, and wing hinges, we found the cosines of direction of the anatomical body axes in the cameras' frame of reference. If $X_B$ is a unit vector in the direction of the longitudinal body axis and $Y_B$ in the direction of the transverse axis (connection line between both wing hinges), then $Z_B$ is defined as the dorsoventral axis by the cross product of $X_B$ and $Y_B$ (Fig. S4A):

$$\widehat{Z_B} = \frac{\widehat{X_B} \times \widehat{Y_B}}{\sin\,\alpha}, \qquad \text{(eq. 3)}$$

at which the cap above the vectors denotes a unit vector and $\alpha$ is the 90° angle between $X_B$ and $Y_B$. The latter angle might slightly vary due to digitization errors. We made certain that all resulting axes were orthogonal by recalculating $Y_B$ based on $\widehat{Z_B}$ and $\widehat{X_B}$:

$$\widehat{Y_B} = \widehat{Z_B} \times \widehat{X_B}. \qquad \text{(eq. 4)}$$

3) The unit vectors of the body axes are then used to form the rotation matrix M, written as:

$$\mathbf{M} = \begin{bmatrix} \widehat{X_B} \\ \widehat{Y_B} \\ \widehat{Z_B} \end{bmatrix} \qquad \text{(eq. 5)}$$

The matrix rotates the position ($P$ ) of each point on the body and wing in the camera frame of reference to the body frame of reference:

$$P_B = M \times P\,, \qquad \text{(eq. 6)}$$

with $P_B$ the desired position in the body frame of reference.

*S2.3 Reconstruction of wing flapping angles*

Wing-flapping angles in the body frame of reference were calculated according to the following procedure (Fig. S2). The elevation angle of the wing (θ) was calculated for each frame in the body's frontal ($Y_BZ_B$) plane as:

$$\theta = \tan^{-1}\left(\frac{WL_Z}{WL_Y}\right) \text{ for left and } \theta = \tan^{-1}\left(\frac{WL_Z}{-WL_Y}\right), \qquad \text{(eqs 7, 8)}$$

and right wing, with $WL$ the wings' longitudinal axis (Fig. S2B) and the subscripts Y and Z denote the relevant components in the body frame of reference. The sweep angle ($\phi$) in the horizontal was calculated for each frame by rotating the wing position data ($P_1$) about the longitudinal axis by $\theta$ so that the wing axis is in the body plane ($X_BY_B$):

$$P_1{}' = \begin{bmatrix} 1 & 0 & 0 \\ 0 & \cos\theta & \sin\theta \\ 0 & -\sin\theta & \cos\theta \end{bmatrix} \times P_1 \,. \qquad \text{(eq. 8)}$$

Sweep angle ($\phi$) is calculated from the rotated data of the left wing as:

$$\phi = \tan^{-1}\left(\frac{WL_X}{WL_Y}\right) \text{ and } \phi = \tan^{-1}\left(\frac{WL_X}{-WL_Y}\right) \qquad \text{(eqs 10, 11)}$$

for the right wing. The angle of incidence angle ($\psi$) is calculated after an additional rotation of the wing data about the insect's dorsoventral axis by $\phi$ to align the wing axis with the $\widehat{Y_B}$-axis. This is achieved by:

$$P_1{}'' = \begin{bmatrix} \cos\phi & -\sin\phi & 0 \\ \sin\phi & \cos\phi & 0 \\ 0 & 0 & 1 \end{bmatrix} \times P_1{}'. \qquad \text{(eq. 12)}$$

The angle of incidence is then:

$$\psi = -\tan^{-1}\left(\frac{WC_Z}{WC_X}\right). \qquad \text{(eq. 13)}$$

The wing's angle of attack (AoA) with respect to the direction of wing motion is calculated from wing length ($WL$) and chord ($WC$) vectors as illustrated in Fig. S2. LV and FV are vectors connecting the wing hinge with leading and trailing edge at 0.7 of the wing length, respectively. The vector (WN) normal to the area of the left wing is calculated from the following cross product:

$$WN = LV \times TV. \qquad \text{(eq. 14)}$$

The vector (WSN) perpendicular to both wing velocity (WV) and the wing length (WL) is:

$$WSN = WV \times WL, \qquad \text{(eq. 16)}$$

and angle of attack is derived as:

$$AoA = \cos^{-1}(WN \cdot WSN). \qquad \text{(eq. 17)}$$

Calculations for the right wing are similar but the order of the cross products are changed according to the right-hand rule. For further explanations, see figure S2. Figure S3 shows kinematics and instantaneous flight forces of 3 additional animals. Movies of air velocity, vorticity and pressure changes of all seven stroke cycles and at t/T = 0, 0.15, 0.35, 0.50, 0.60, 0.70, 0.95 are available by request from the authors.

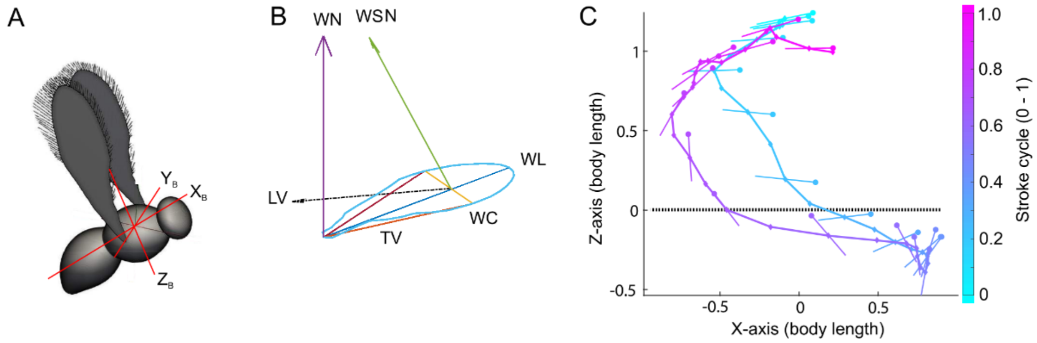


**Figure S2.** Definitions and reconstruction of body and wing kinematics. **(**A) Boy frame of reference of the coordinate system. (B) Coordinate system of the wing used to find the angle of attack (AoA). $X_W$ is wing length between tip and hinge, $Y_W$ is wing chord between leading and trailing edges at 0.7 wing length, and $Z_W$ is a cross-product of vectors pointing towards the leading (LV) and trailing (TV) edges. Wing length (WL) is denoted in blue, wing chord (WC) in yellow, LV in red and TV in orange. WN is the vector normal to the wing area (pointing towards the wing's dorsal side. WV is the wing velocity vector (shown as a dashed black arrow). WSN (green) is a vector that is normal to both the wing length and wing velocity. (C) Example of the raw wing kinematic data extracted from the high-speed videos. The above figure is shown as figure 7A in the main text after smoothing by Fourier filtering restricted to five modes.

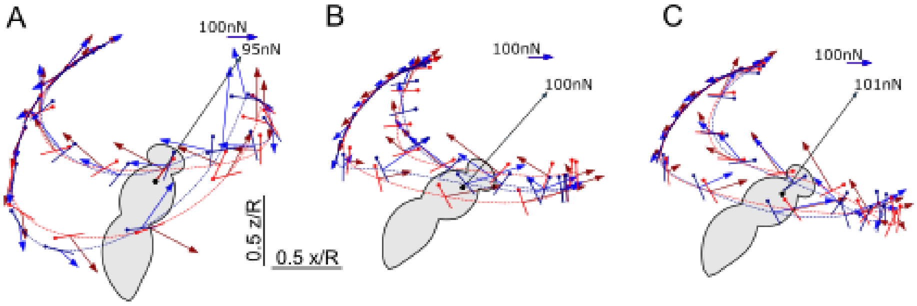


**Figure S3.** Instantaneous force vectors from the CFD overlaid on the simulated wingtip trajectory of 3 additional wasps. The simulated trajectories are based on the observed kinematics after Fourier smoothing. **(**A-B) Kinematics and forces of wasp 2*1 (*A*, horizontal velocity, 0.145 ms$^{-1}$), 3*2 (*B*, horizontal velocity, 0.172 ms$^{-1}$), and 3*1 (*C*, horizontal velocity, 0.195 ms$^{-1}$, Table 1). Dashed lines denote wing tip trajectory, colour-coded solid line denote the wing's chord where the leading edge is marked by a circle. Arrows denote instantaneous force vectors whereas the single black arrow denoted the (time) average force for the cycle. Blue and red colours denote the right and left wings, respectively.

##### *S2.4 Parameters of body dynamics*

Besides wing motion, our CFD modelling also considers the changes in body posture angles about the three body axes (yaw, pitch, roll) within the stroke cycle. We approximated the changes in posture within the stroke cycle by a first order Fourier approximation using sine and cosine fit components (cf. Fig. 3 of the main text). These parameters are shown in Table S1.

**Table S1.** Parameters of body dynamics that were used for numerical modelling.

| | Mean (zero coefficient) | | | 1st cosine harmonic | | | 1st sine harmonic | | |
|---|---|---|---|---|---|---|---|---|---|
| Animal*stroke | Yaw (°) | Pitch (°) | Roll (°) | Yaw (°) | Pitch (°) | Roll (°) | Yaw (°) | Pitch (°) | Roll (°) |
| 1*1 | 60.8 | -36.7 | 15.8 | -0.66 | -1.72 | -1.51 | -4.63 | -2.86 | 8.14 |
| 1*2 | 41.2 | -28.7 | -2.5 | -4.79 | -5.00 | 2.07 | 2.27 | -1.24 | -0.99 |
| 2*1 | 32.3 | -65.2 | 8.9 | -1.55 | -4.64 | -1.92 | 1.65 | -1.05 | 0.06 |
| 2*2 | 31.9 | -67.1 | 11.4 | -1.55 | -4.64 | -1.92 | 1.65 | -1.05 | 0.06 |
| 2*3 | 40.1 | -67.3 | -1.9 | 3.00 | -3.14 | -1.84 | 5.50 | 0.00 | -6.64 |
| 3*1 | 16.3 | -35.1 | -23.6 | 2.25 | -3.61 | -7.96 | -7.04 | -0.78 | 7.80 |
| 3*2 | 10.1 | -35.2 | 0.1 | -2.29 | -4.53 | -1.97 | -4.50 | 0.32 | 5.29 |

*S2.5 Inertial power requirements for wing flapping*

In addition to calculations of inertial power for wing flapping using Ellington's framework [1] based on mean values, we also derived instantaneous inertial power for the tested animals. In contrast to Ellington, we here use the measured wing kinematics and a more elaborated estimation of the wing's centre of mass. We assumed that (1) membranous wing mass is 0.076 µg (1% of the total body mass [2]), (2) The bristles contribute to the inertia, and 3) wing thickness is uniform. From these data, we calculated mass moments of inertia of $I_{xx}$=12049 µg µm$^2$, $I_{yy}$=208 µg µm$^2$ and $I_{zz}$=12257 µg µm$^2$ for the forewing and $I_{xx}$=2020 µg µm$^2$, $I_{yy}$=149 µg µm$^2$ and $I_{zz}$=2169 µg µm$^2$ for the hindwing. To compare, values reported for the fully bristled and slightly smaller wing of *Paratuposa placentis* were 3370 µg µm$^2$, 360 µg µm$^2$ and 3730 µg µm$^2$, respectively. Figure S4 shows instantaneous total inertial power, aerodynamic power, and the sum of inertial and aerodynamic power without elastic storage for flapping both wings (cf. tables 1 and 2). Table S2 shows mean inertial power, aerodynamic power and total flight power assuming both perfect elastic energy storage and no elastic energy storage.

**Table S2.** Mean power for wing flapping averaged over one stroke cycle. $P_{acc}$, inertial power; $P_{aero}$, aerodynamic power; $P_{total}$, sum of inertial and aerodynamic power at 0% and 100% elastic energy storage (e.s.).

| Animal*stroke | $P_{acc}$ (nW) | $P_{aero}$ (nW) | $P_{total}$ 0% e.s. (nW) | $P_{total}$ 100% e.s. (nW) |
|---|---|---|---|---|
| 1*1 | -0.6 | 235.6 | 251.1 | 235.0 |
| 1*2 | -1.7 | 289.8 | 298.3 | 288.0 |
| 2*1 | -1.1 | 279.2 | 286.6 | 278.1 |
| 2*2 | -1.3 | 251.6 | 258.8 | 250.2 |
| 2*3 | -1.3 | 263.3 | 271.1 | 262.0 |
| 3*1 | 0.18 | 266.5 | 269.4 | 266.7 |
| 3*2 | -1.3 | 292.5 | 296.7 | 291.2 |

*S2.6 Grid independence study*

To verify that the results are independent of the resolution of the numerical simulations, we performed a grid independence study for each simulation presented in the main text. Figure S4 shows the vertical force for the simulation 3*2 (Animal*cycle). The resolution differs by a factor of two, the coarse resolution being $\Delta x$/R =512 and $\Delta x/h_w$=4.2, and the high resolution twice that values. The largest instantaneous difference in the vertical force is only 2.76%, indicating that the results are indeed independent of $\Delta x$. The coarser resolution is thus used for all results in the main text

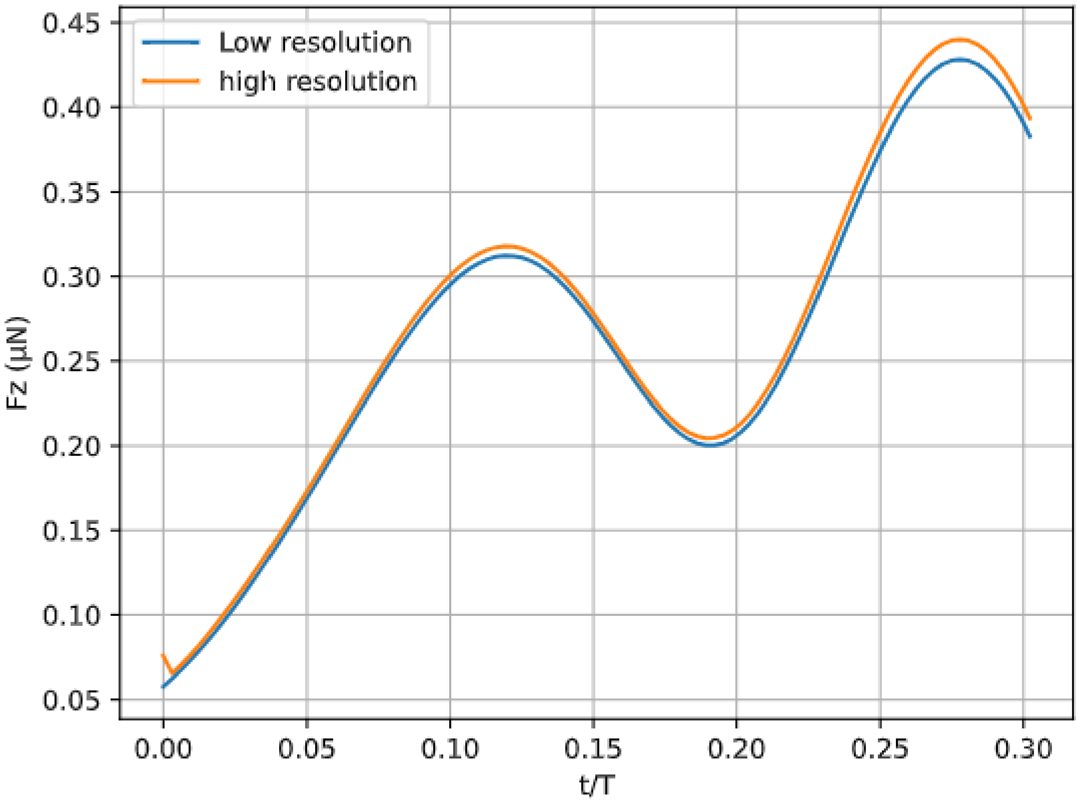


**Figure S4.** Grid independence study. Shown are the vertical forces (Fz) simulated for cycle 3*2 with a grid resolution of $\Delta x$/R =512 and $\Delta x/h_w$=4.2 (blue line) and the result simulated at twice higher resolution (orange)

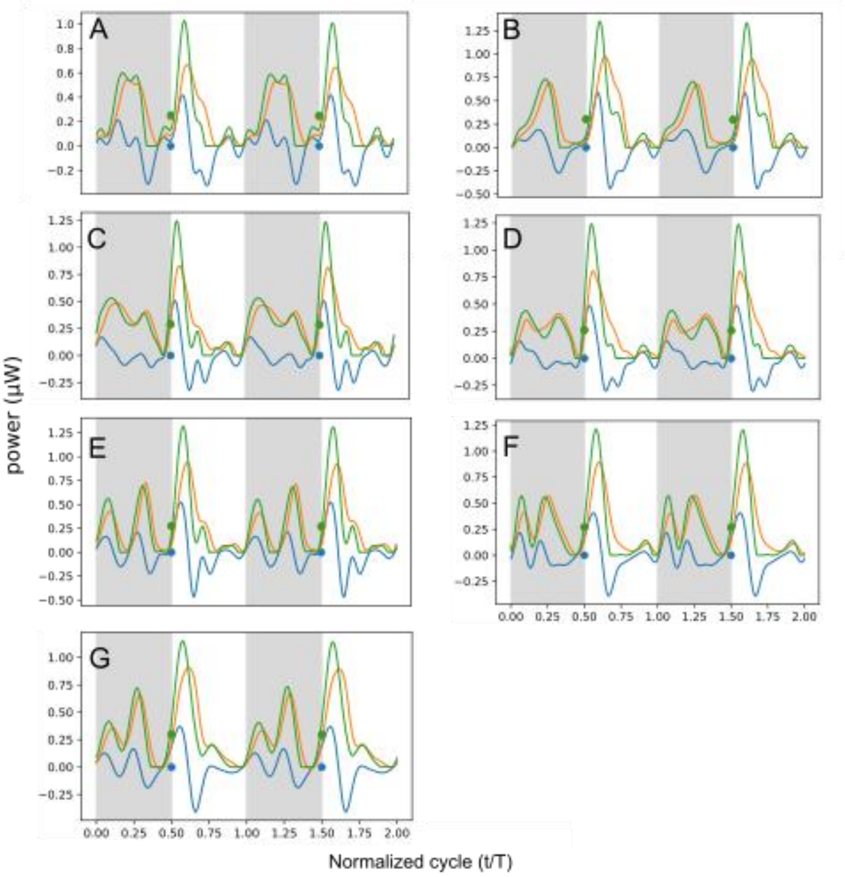


**Figure S5.** Instantaneous power for wing flapping. Traces show inertial power for both wings (blue), aerodynamic power (orange), and the sum of inertial and aerodynamic power without elastic energy storage (green). Animal and stroke cycle (cf. table 1) are (A) 1*1, (B) 1*2, (C) 2*1, (D) 2*2, (E) 2*3, (F) 3*1, (G) 3*2.

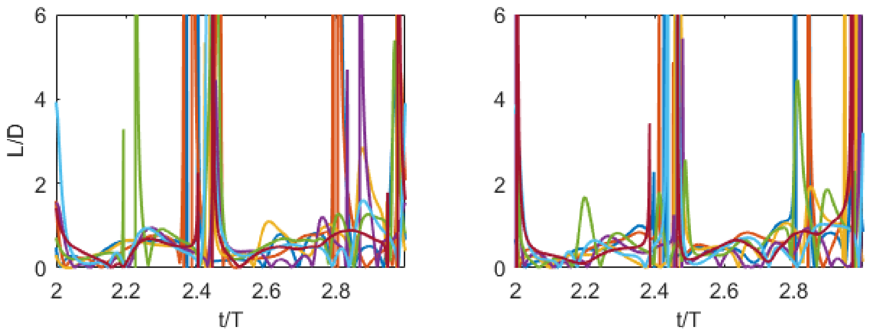


**Figure S6.** Lift/drag force ratios for the wing profile. Lines depict the ratios of computed instantaneous wing lift and drag generated by the moving left and right forewings (left and right plots respectively) during a flapping cycle. Line colours denote the seven different flapping cycles.

**References**

[1] Ellington, C. P. (1984). The aerodynamics of hovering insect flight. IV. Lift and power requirements. *Phil. Trans. Roy. Soc. Lond.* B **305**(1122), 145-181.

[2] Farisenkov, S.E., Kolomenskiy, D., Petrov, P.N., Engels, T., Lapina, N.A., Lehmann, F.-O., Onishi, R., Liu, H. & Polilov, A.A. 2022 Novel flight style and light wings boost flight performance of tiny beetles. *Nature* **602**, 96-100.
